# Gene identification for ocular congenital cranial motor neuron disorders using human sequencing, zebrafish screening, and protein binding microarrays

**DOI:** 10.1101/2024.09.12.612713

**Authors:** Julie A. Jurgens, Paola M. Matos Ruiz, Jessica King, Emma E. Foster, Lindsay Berube, Wai-Man Chan, Brenda J. Barry, Raehoon Jeong, Elisabeth Rothman, Mary C. Whitman, Sarah MacKinnon, Cristina Rivera-Quiles, Brandon M. Pratt, Teresa Easterbrooks, Fiona M. Mensching, Silvio Alessandro Di Gioia, Lynn Pais, Eleina M. England, Teresa de Berardinis, Adriano Magli, Feray Koc, Kazuhide Asakawa, Koichi Kawakami, Anne O’Donnell-Luria, David G. Hunter, Caroline D. Robson, Martha L. Bulyk, Elizabeth C. Engle

**Affiliations:** F.M. Kirby Neurobiology Center, Boston Children’s Hospital, Boston, MA, USA; Department of Neurology, Boston Children’s Hospital, Boston, MA, USA; Department of Neurology, Harvard Medical School, Boston, MA, USA; Broad Institute of MIT and Harvard, Cambridge, MA, USA; Division of Genetics, Department of Medicine, Brigham and Women’s Hospital and Harvard Medical School, Boston, MA, USA; Howard Hughes Medical Institute, Chevy Chase, MD, USA; Bioinformatics and Integrative Genomics Graduate Program, Harvard University, Cambridge, MA 02138, USA; Department of Ophthalmology, Boston Children’s Hospital, Boston, MA, USA; Department of Ophthalmology, Harvard Medical School, Boston, MA, USA; Regeneron Pharmaceuticals, Tarrytown, NY, USA; Program in Medical and Population Genetics, Broad Institute of MIT and Harvard, Cambridge, MA, USA; Division of Genetics and Genomics, Boston Children’s Hospital, Harvard Medical School, Boston, MA, USA; Department of Ophthalmologic Sciences, Faculty of Medicine and Surgery, University “Federico II”, Naples, Italy; Department of Ophthalmology, Faculty of Medicine, Izmir Katip Celebi University, Izmır, Turkey; Neurobiology and Pathology Laboratory, National Institute of Genetics, Mishima, Shizuoka, Japan; Laboratory of Molecular and Developmental Biology, National Institute of Genetics; Department of Genetics, Graduate University for Advanced Studies (SOKENDAI); Center for Genomic Medicine, Massachusetts General Hospital, Boston, MA, USA; Division of Neuroradiology, Department of Radiology, Boston Children’s Hospital, Boston, MA, USA; Department of Radiology, Harvard Medical School, Boston, MA, USA; Department of Pathology, Brigham and Women’s Hospital and Harvard Medical School, Boston, MA, USA

## Abstract

**Purpose:** To functionally evaluate novel human sequence-derived candidate genes and variants for unsolved ocular congenital cranial dysinnervation disorders (oCCDDs).

**Methods:** Through exome and genome sequencing of a genetically unsolved human oCCDD cohort, we previously identified variants in 80 strong candidate genes. Here, we further prioritized a subset of these (43 human genes, 57 zebrafish genes) using a G0 CRISPR/Cas9-based knockout assay in zebrafish and generated F2 germline mutants for seventeen. We tested the functionality of variants of uncertain significance in known and novel candidate transcription factor-encoding genes through protein binding microarrays.

**Results:** We first demonstrated the feasibility of the G0 screen by targeting known oCCDD genes *phox2a* and *mafba*. 70-90% of gene-targeted G0 zebrafish embryos recapitulated germline homozygous null-equivalent phenotypes. Using this approach, we then identified three novel candidate oCCDD genes (*SEMA3F*, *OLIG2,* and *FRMD4B*) with putative contributions to human and zebrafish cranial motor development. In addition, protein binding microarrays demonstrated reduced or abolished DNA binding of human variants of uncertain significance in known and novel sequence-derived transcription factors *PHOX2A* (p.(Trp137Cys)), *MAFB* (p.(Glu223Lys)), and *OLIG2* (p.(Arg156Leu)).

**Conclusions:** This study nominates three strong novel candidate oCCDD genes (*SEMA3F*, *OLIG2,* and *FRMD4B*) and supports the functionality and putative pathogenicity of transcription factor candidate variants *PHOX2A* p.(Trp137Cys), *MAFB* p.(Glu223Lys), and *OLIG2* p.(Arg156Leu). Our findings support that G0 loss-of-function screening in zebrafish can be coupled with human sequence analysis and protein binding microarrays to aid in prioritizing oCCDD candidate genes/variants.

## Introduction

Human exome and genome sequencing have facilitated Mendelian gene discovery but generate numerous candidate genes and variants, only a subset of which are pathogenic. Moreover, heterogeneous loci and alleles hinder the identification of recurrently mutated genes or alleles. Targeted modeling can assess functionality but is often limited in throughput and applicability across disparate genes and phenotypes. This disconnect between candidate disease gene identification and functional validation remains a barrier to gene discovery for phenotypes including the ocular congenital cranial dysinnervation disorders (oCCDDs).

oCCDDs are characterized by congenitally restricted eye and/or eyelid movement and result from maldevelopment of the oculomotor (CN3), trochlear (CN4), or abducens (CN6) motor neurons and/or axons (Fig. 1A-B). Some oCCDDs include congenital ptosis, Marcus Gunn jaw-winking syndrome (MGJWS), congenital fibrosis of the extraocular muscles (CFEOM), and Duane retraction syndrome (DRS). Congenital ptosis is characterized by eyelid drooping and can result from CN3 maldevelopment. In MGJWS, congenital ptosis transiently improves with specific jaw movements due to synkinetic miswiring by the motor trigeminal nerve (CN5). CFEOM results from maldevelopment of CN3 and in some cases also CN4 or CN6, and is typified by non-progressive restriction of vertical eye movement with variable ptosis and variable restrictions of horizontal gaze. In DRS, CN6 maldevelopment causes limited abduction and variably limited adduction, and the globe retracts upon attempted adduction due to synkinetic miswiring by CN3.

**Figure 1.**
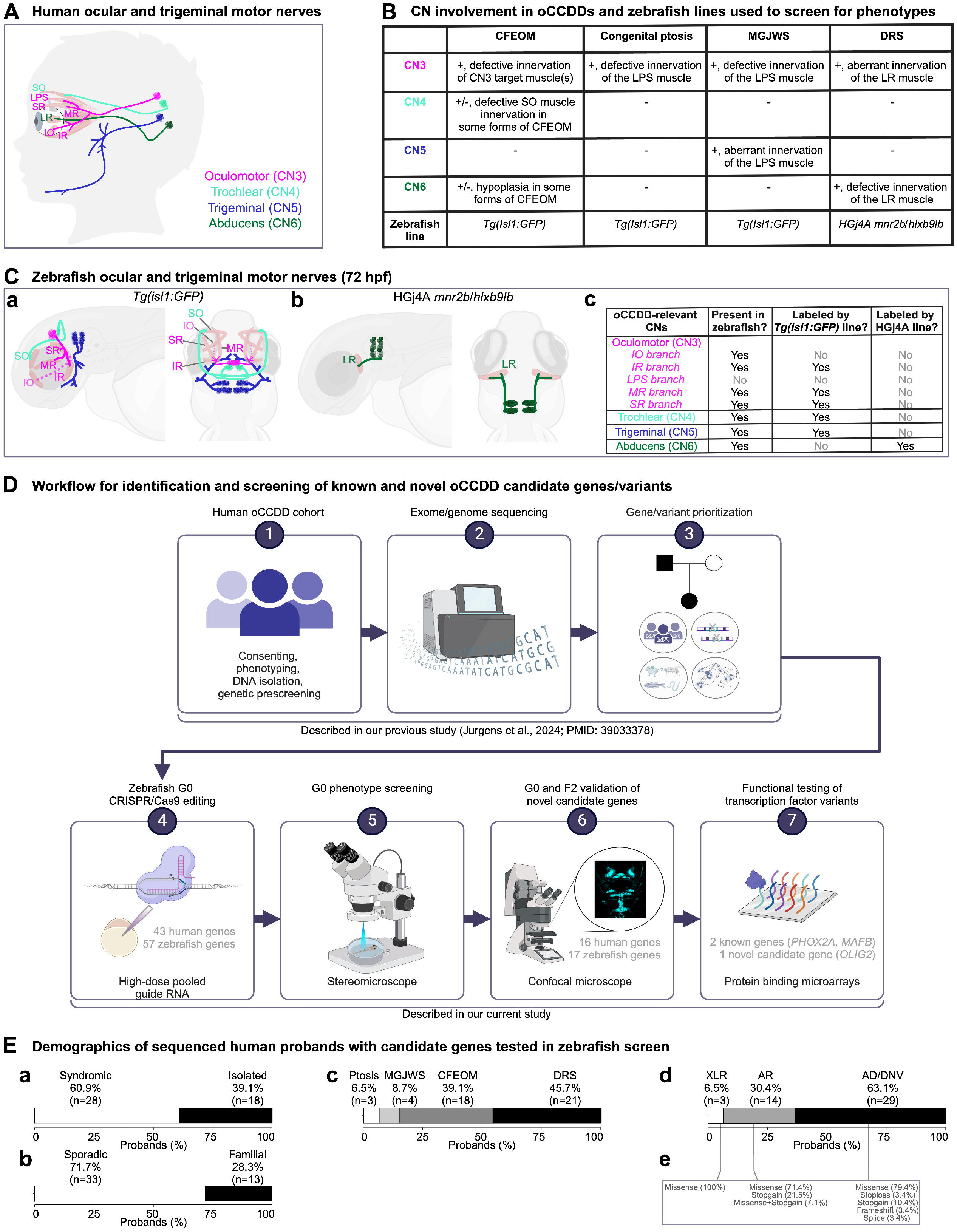
Human and zebrafish cranial motor nerves and study workflow. **A:** In the human brainstem, motor neurons cluster into cranial motor nuclei, whose axons form the cranial motor nerves (CNs). oCCDDs can arise from defective formation, identity, and/or axonal projections of three of these motor neuron populations (CN3, CN4, and CN6) that collectively innervate seven extraocular muscles to orchestrate eye/ eyelid movement. CN3 innervates the IO, IR, LPS, MR, and SR muscles. CN4 innervates the SO muscle, and CN6 innervates the LR muscle. CN5 normally innervates the muscles of mastication (not shown) but can also aberrantly innervate the LPS muscle in MGJWS. **B:** Summary of CN pathology in humans or mice with specific oCCDDs, and the zebrafish reporter lines used to analyze them. To model oCCDDs in zebrafish, our study leveraged the HGj4A *mnr2b*/*hlxb9lb* line for DRS candidate genes and the *Tg(isl1:GFP)* line for CFEOM, congenital ptosis, and MGJWS genes. **C:** Zebrafish wild-type ocular and trigeminal motor nerves visualized in the *Tg(isl1:GFP)* (a) or HGj4A reporter line (b). Left-lateral view; right-dorsal view. c) Summary of oCCDD-relevant CNs that are present in zebrafish and labeled by each reporter line. (a,c): As in humans, zebrafish CN3 innervates the IO, IR, MR, and SR muscles. However, the CN3 branch to the IO muscle cannot be visualized with the *Tg(isl1:GFP)* transgenic line. Additionally, unlike humans, zebrafish lack the LPS muscle and its corresponding CN3 subdivision. Zebrafish have bilateral CN4 motor nuclei and nerves (left and right). CN4 exits the brainstem dorsally and travels ventrally to innervate the contralateral SO muscle. At 72 hpf, wild-type zebrafish CN4 is variably defasciculated. Zebrafish have anterior and posterior trigeminal motor nuclei bilaterally, which extend axons for the motor trigeminal (CN5) nerve that innervates the muscles of mastication (not shown). (b,c): The HGj4A line labels the CN6 motor neurons and their axons. Zebrafish have anterior and posterior CN6 motor nuclei bilaterally, which extend CN6 nerves that target the LR muscles. **D:** Workflow for the identification and screening of known and novel oCCDD candidate genes and variants. We reported steps 1-3 in our previous study describing exome/ genome sequencing of our human oCCDD cohort^3^ and performed steps 4-7 in the present study. **E:** Demographics of the 46 human probands whose candidate genes were tested in the zebrafish screen. Percentages of probands are shown with oCCDDs that are: a) syndromic or isolated, b) sporadic or familial, c) fitting various oCCDD subdiagnoses, or d) fitting various modes of inheritance. e) The types of variants fitting modes of inheritance defined in d) are provided. Abbreviations: AD-autosomal dominant, AR-autosomal recessive, CFEOM-congenital fibrosis of the extraocular muscles, CN-cranial nerve, CN3-oculomotor/ cranial nerve 3, CN4-trochlear/ cranial nerve 4, CN5-trigeminal motor/ cranial nerve 5, CN6-abducens/ cranial nerve 6, DNV-de novo variant, DRS-Duane retraction syndrome, hpf-hours post-fertilization, IO-inferior oblique muscle, IR-inferior rectus muscle, LPS-levator palpebrae superioris muscle, LR-lateral rectus muscle, MGJWS-Marcus Gunn jaw-winking syndrome, MR-medial rectus muscle, oCCDD-ocular congenital cranial dysinnervation disorder, Ptosis-congenital ptosis, SO-superior oblique muscle, SR-superior rectus muscle, XLR-X-linked recessive. Key: pink-CN3, light green-CN4, dark blue-CN5, dark green-CN6, partially transparent brown-extraocular muscle, dashed line-nerve that is present in zebrafish but not labeled with the transgenic reporter line.

Some oCCDDs are caused by monoallelic or biallelic loss of function (LOF) of transcription factors. Biallelic *PHOX2A* LOF causes CFEOM in humans and absence of CN3/CN4 motor nuclei in mice.^1^ Monoallelic *MAFB* LOF causes DRS in humans and absence of CN6 motor nuclei in mice, with secondary aberrant innervation by CN3 of the lateral rectus muscle, which is normally innervated by CN6.^2^ While these and other mechanisms explain some oCCDDs, many remain genetically undefined. We recently reported exome/genome sequencing of a large human oCCDD cohort, which yielded many candidate genes and variants of uncertain significance whose pathogenicity remain untested.^3^

Targeted *in vitro* or *in vivo* modeling can enhance functional understanding of these candidate genes/ variants. For instance, candidate variants in transcription factors can be assessed by universal protein binding microarrays, a high-throughput method that tests DNA binding capacity of wild-type or variant transcription factors.^4–5^ Additionally, oCCDD genetics and pathophysiology can be elucidated with animal models including zebrafish (*Danio rerio*). Zebrafish have advantages over mammals, including fecundity and rapid external development.^6^ Moreover, 69% of human genes and 82% of human disease-associated genes in Online Mendelian Inheritance in Man (OMIM)^7^ have fish orthologs; some of these human genes are represented as duplicated paralogous genes in fish.^8^ Zebrafish are amenable to gene editing and live imaging, and their ocular motor neurons are detected by 24-30 hours post-fertilization (hpf) and innervate their target muscles by 72 hpf.^9^ Zebrafish lack eyelids and the CN3 branch that innervates them, but their cranial nerve anatomy and eye movements are otherwise conserved.^6,9^

Zebrafish ocular motor development has been characterized through transgenic reporters including a *Tg*(*isl1:GFP*) line labeling all cranial motor neurons and their axons except CN6,^10^ and an HGj4A *mnr2b*/*hlxb9lb* enhancer trap line labeling CN6 motor neurons and axons (Fig. 1C).^11^ As in mammals, *phox2a^-/-^*fish have absent or malformed CN3 and CN4 nuclei,^12–14^ and *mafba^-/-^* fish have absent or hypoplastic CN6 nuclei.^15–17^ Thus, zebrafish are an excellent model for moderate-throughput testing of oCCDD candidate genes.

Standard homozygous null zebrafish are derived in the F2 generation and require 6 months to generate. Prior studies have expedited this timeline by generating G0 mutants using simultaneous injection of multiple high-dose CRISPR guide RNAs redundantly targeting one gene.^18–21^ This approach is reported to reproduce homozygous null-equivalent phenotypes in >90% of embryos with <18% toxicity. Since the cranial motor system is conserved between zebrafish and humans and there are existing transgenic lines labeling the ocular cranial motor system, oCCDDs represent a unique model for testing the G0 knockout approach.

Here we report the successes and limitations of functionally assessing human candidate oCCDD genes using moderate-throughput CRISPR/Cas9 G0 LOF screening in zebrafish, and testing functionality of transcription factor candidate variants by protein binding microarrays (Fig. 1D).

## Methods

Additional details for the following sections are provided in Supplementary Methods.

### Human consenting and phenotyping

Our study was approved by the Boston Children’s Hospital (BCH) Institutional Review Board (IRB 05-03-036R) and complied with ethical recommendations of BCH and the Declaration of Helsinki. Research participants or legal guardians provided written informed consent. Phenotypes were obtained from review of clinical records and from participant questionnaires and updates. Clinically acquired brain magnetic resonance imaging exams (MRIs) were reviewed retrospectively.

### Prioritization and cosegregation analysis of human sequence-derived alleles for screening in zebrafish

In our previous study, we leveraged human genetics to identify novel oCCDD candidate genes/variants through phenotyping and exome/genome sequencing of a large cohort of human pedigrees with oCCDDs.^3^ In the present study, we further prioritized these human sequence-derived candidates for the zebrafish screen based on conservation in zebrafish, recessive inheritance, and/or predicted LOF of variants (Fig. 1D, Supplementary Table 1). Sanger validation and familial cosegregation analysis are shown for candidate variants in genes that 1) yielded cranial motor phenotypes in both G0 and F2 mutants in our zebrafish screen, and/or 2) encoded transcription factors tested by protein binding microarray (Supplementary Table 2).

### G0 screening and F2 germline validation in LOF zebrafish models of prioritized genes

Zebrafish studies were conducted in accordance with the *ARVO* Statement for the Use of Animals in Ophthalmic and Vision Research. G0 targeting experiments consisted of microinjecting single cell-stage embryos with four high dose (1 ug/uL) guide RNAs redundantly targeting each gene.^20^ At 72 hpf, injected G0 fish were assessed by stereomicroscope for gross phenotypic changes in cranial motor neuron nuclei and/or nerves. For genes whose targeting induced putative phenotypes in at least a subset of injected fish, we performed two additional G0 experimental replicates and F2 germline mutant validation and visualized with confocal imaging (Fig. 1D).

### Protein structural mapping and universal protein binding microarray testing of transcription factor candidate variants

2D protein structural maps were generated for genes/variants validated through the zebrafish screen and/or protein binding microarrays. Protein binding microarrays were used to assess DNA binding capabilities of variants of uncertain significance in the DNA binding domains of known (*PHOX2A, MAFB*) or novel (*OLIG2*) transcription factor-encoding candidate genes relative to their wild-type counterparts (Fig. 1D).^5,22^

## Results

### Prioritization of human sequence-derived oCCDD candidate genes for screening in zebrafish

For G0 LOF targeting and anatomic screening in zebrafish, we chose 43 human candidate genes from our previously reported analysis of exome/genome sequences of 467 oCCDD pedigrees (Fig. 1D).^3^ Three of the 43 genes had rare variants of interest in two probands (*ACTR1B, KCNAB1, FOXC1;* Fig. 1D, Supplementary Table 1), two had well established roles in oCCDDs before this study (*PHOX2A, MAFB*), and four had only occasional reported association with human oCCDDs (*ARX, COL25A1, DYRK1A, KIFBP*). Thirty-seven were novel oCCDD candidates, seven of which were highlighted in our previous report.^3^

Summaries of the oCCDDs, modes of inheritance, and nature of the candidate variants in these 46 probands are provided (Fig. 1E). Probands predominantly had syndromic (60.9%) and sporadic (71.7%) oCCDDs, most of which were DRS (45.7%) or CFEOM (39.1%). Most candidate variants were identified under autosomal dominant/ *de novo* models (63.1%), and most were missense (78.3% across all inheritance models) with fewer putative LOF variants (21.7%).

The zebrafish orthologs of these 43 human genes were subjected to a G0 CRISPR/Cas9 editing pipeline in zebrafish. Fourteen of the human genes had two highly homologous zebrafish paralogs (*CELF5, DBX1, DYRK1A, FOXC1, FRMD4B, MXRA8, NAV2, NTN1, PPP1R14B, SDK1, SEMA3F, SEMA5B, SLC12A5, TLE3*), and thus fifty-seven zebrafish genes were targeted. Each zebrafish paralog was targeted independently.

### G0 screen and F2 zebrafish modeling for human sequence-derived oCCDD candidate genes

To pilot the G0 LOF zebrafish assay, we targeted the known oCCDD genes *phox2a* and *mafba* (Fig. 2A-H; Fig. 3A-J). G0 targeting of both genes recapitulated known germline LOF phenotypes. G0-targeted fish had grossly absent (*phox2a*-69.0%, *mafba*-78.9%) or malformed (*phox2a*-10.9%, *mafba*-8.6%) CN3/CN4 or CN6 motor neuron nuclei, respectively, with absence or thinning of the corresponding cranial nerves (Fig. 2A-G, Fig. 3A-I). Targeting of each gene had no significant impact on survival (Fig. 2H, Fig. 3J).

**Figure 2.**
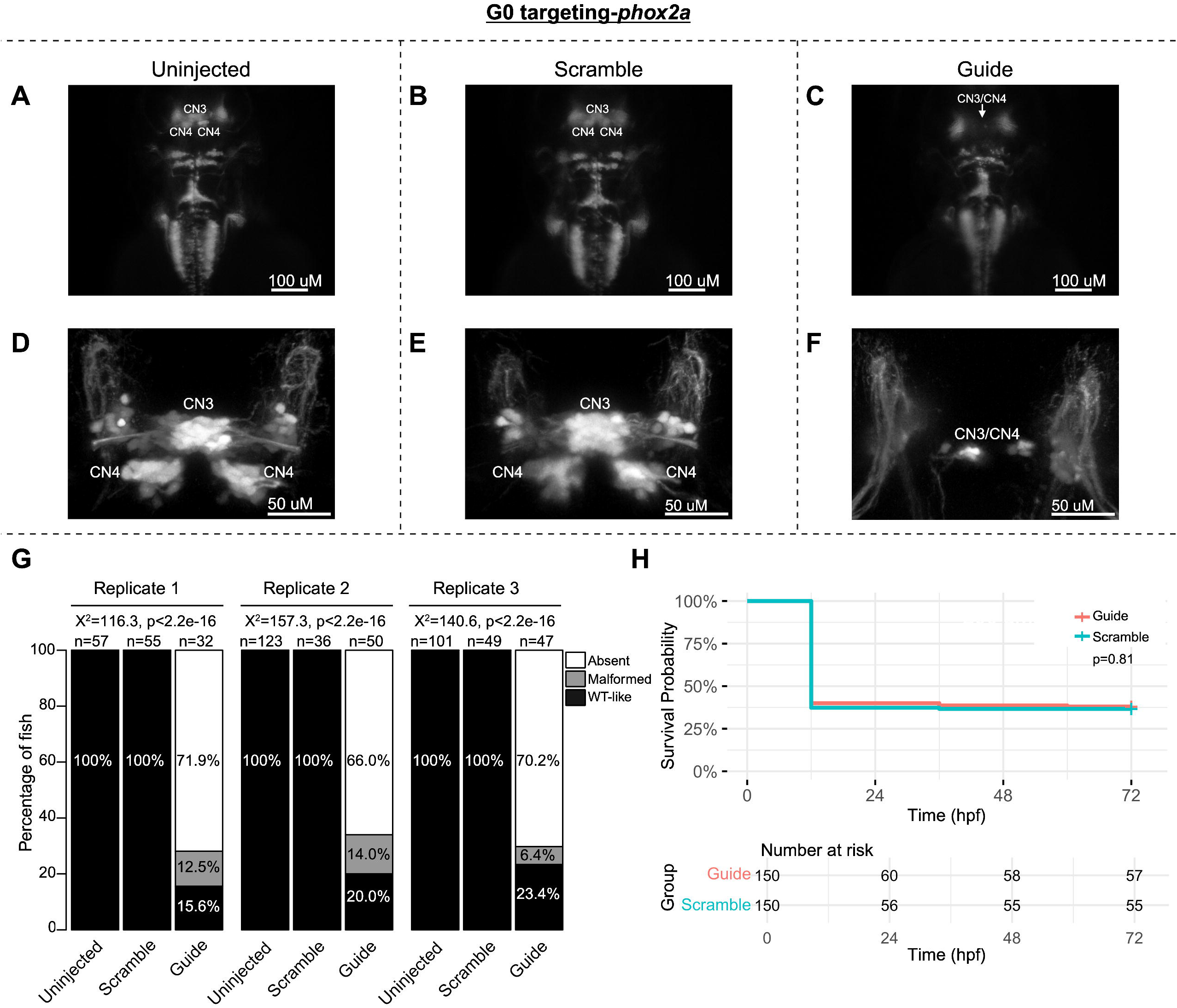
G0 zebrafish targeting of *phox2a*. Representative images obtained by stereomicroscope **(A-C)** and confocal microscopy **(D-F)** of Uninjected (A,D), Scramble-injected (B,E), or *phox2a*-targeting Guide-injected G0 *Tg(isl1:GFP)* zebrafish at 72 hpf. Stereomicroscope images for each treatment group correspond to the confocal images obtained from the same fish in the panel below. Since residual motor neurons from CN3 and/or CN4 could not be definitively assigned to either of these specific motor neuron nuclei, CN3/CN4 were labeled as a single entity in Guide-targeted fish. Note apparent absence (C) or paucity (F) of CN3/CN4 motor neurons following *phox2a* G0 mosaic knockout. **G:** Barplots showing the percentages of zebrafish exhibiting wild-type-like (“WT-like”), malformed, or absent CN3/CN4 motor nuclei, as scored under the dissecting stereomicroscope in G0 zebrafish at 72 hpf. Total numbers (n’s) of fish in each group are given above the corresponding bar. Data are shown from three experimental replicates. Pearson’s 3x3 chi-squared test with 4 degrees of freedom; p-values and chi-squared values provided for each replicate. **H:** Kaplan-Meier survival curves demonstrating the relative survival probabilities of Scramble-injected (blue line) and Guide-injected (pink line) zebrafish over the first 72 hours of life. “Number at risk” below the plot provides the counts of surviving embryos in each group taken every 24 hpf over a 72 hpf period. Relative survival probabilities of *phox2a*-targeting and scrambled gRNA-injected embryos were compared by the log-rank test. Displayed data were derived from a single experimental replicate, but measurements were taken for three experimental replicates, all of which showed the same trend.

**Figure 3.**
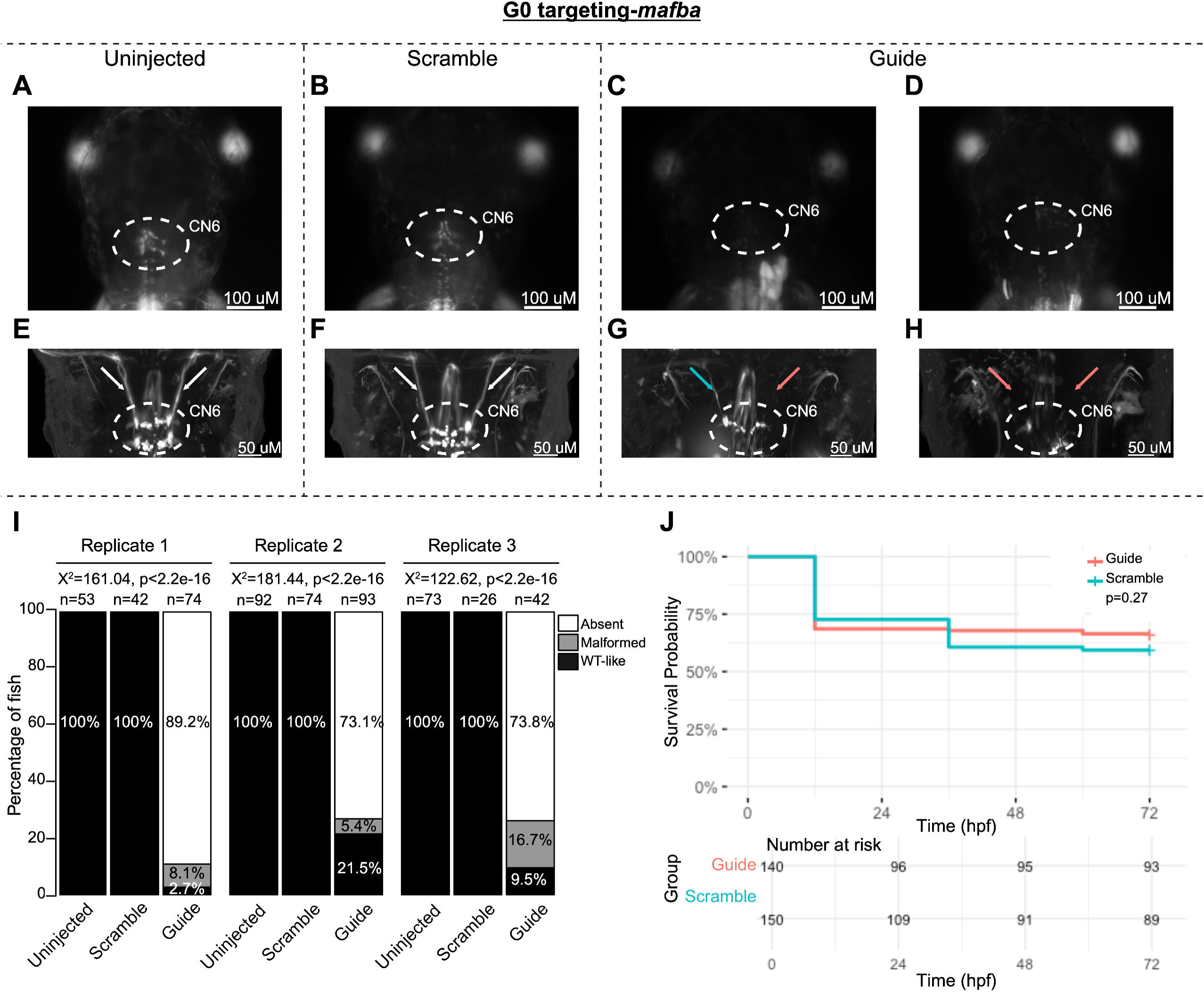
G0 zebrafish targeting of *mafba*. **A-H:** Representative images obtained by stereomicroscope **(A-D)** and confocal microscopy **(E-H)** of Uninjected (A, E), Scrambled guide injected (B, F), or *mafba*-targeting Guide-injected (C-D, G-H) G0 HGj4A zebrafish at 72 hpf. Two images are provided for Guide-injected fish, demonstrating variability in G0 targeting outcomes. Stereomicroscope images for each treatment group correspond to the confocal images obtained from the same fish in the panel below. Motor neurons in the abducens nucleus (dashed white circled region) appear absent by stereomicroscopy (C, D) or variably reduced (G, H) by confocal microscopy, and abducens nerves appear thin (G, blue arrow) or absent (G, H, red arrows) compared to normal (E, F; white arrows) by confocal microscopy. **I:** Barplots showing the percentages of zebrafish exhibiting wild-type-like (“WT-like”), malformed, or absent CN6 motor neuron nuclei, as scored under the dissecting stereomicroscope in G0 zebrafish at 72 hpf. Total numbers (n’s) of fish in each group are given above the corresponding bar. Data are shown from three experimental replicates. Pearson’s 3x3 chi-squared test with 4 degrees of freedom; p-values and chi-squared values provided for each replicate. **J:** Kaplan-Meier survival curves demonstrating the relative survival probabilities of Scramble-injected (blue line) and Guide-injected (pink line) zebrafish over the first 72 hours of life. “Number at risk” below the plot provides the counts of surviving embryos in each group taken every 24 hpf over a 72 hpf period. Relative survival probabilities of *mafba*-targeting Guide-injected and Scrambled-injected embryos were compared by the log-rank test. Displayed data were derived from a single experimental replicate, but measurements were taken for three experimental replicates, all of which showed the same trend.

Encouraged by these results, we performed G0 targeting of the remaining 55 zebrafish oCCDD candidate genes (41 human genes; Supplementary Table 1). These included 32 zebrafish candidate genes (23 human genes, 9 of which were duplicated in fish) for CFEOM, ptosis, or MGJWS and 23 zebrafish candidate genes (18 human genes, 5 of which were duplicated in fish) for DRS. In preliminary results from one pilot injection experiment per gene, G0 mutants for 17 novel zebrafish candidate oCCDD genes (16 human genes) appeared to have at least mild malformation of cranial motor nuclei/ nerves in at least a subset of G0 embryos visualized by stereomicroscope (Supplementary Table 4). The ocular cranial nerve anatomy of these 17 fish were then examined at higher resolution with confocal microscopy of additional G0 targeting replicates and F2 germline mutants. The oCCDD phenotype was confirmed in three putative novel oCCDD candidate genes: *sema3fa, olig2,* and *frmd4bb* (Fig. 4-6, respectively; Supplementary Figures 1-2).

**Figure 4.**
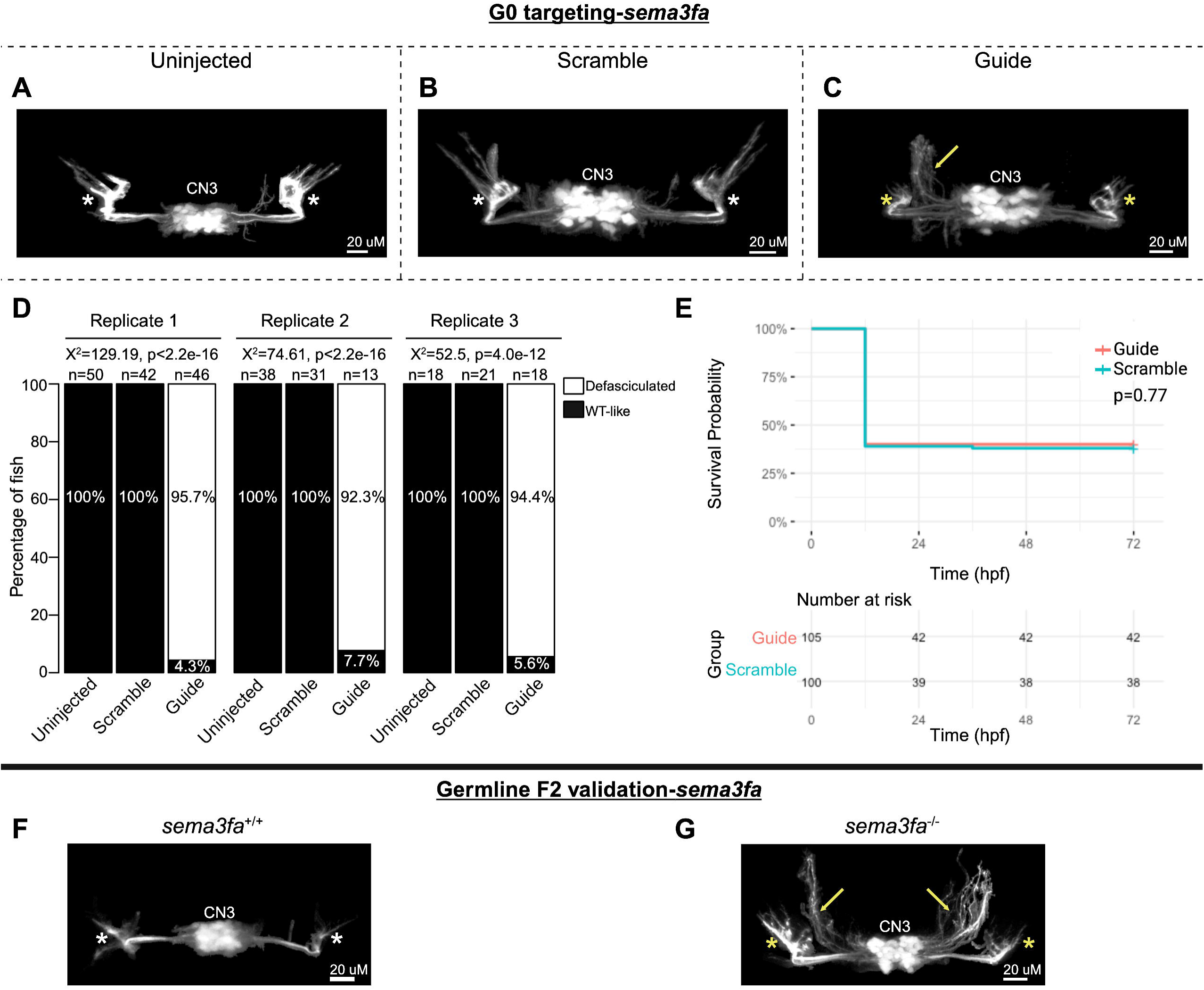
G0 mosaic and F2 germline zebrafish targeting of *sema3fa*. **A-E:** Initial *sema3fa* targeting experiments were performed in G0 embryos. Representative images obtained by confocal microscopy **(A-C)** of Uninjected (A), Scrambled guide injected (B), or *sema3fa*-targeting Guide-injected (C) G0 *Tg(isl1:GFP)* zebrafish at 72 hpf. Note defasciculated CN3 (yellow arrow) and failed CN3 nerve extension toward extraocular muscles (yellow stars) relative to Uninjected and Scramble-injected CN3 (white stars). Key: CN3-cranial nerve 3 (oculomotor), yellow arrow-increased defasciculation of CN3 nerve, yellow star-failed extension of CN3 nerve toward target extraocular muscles, white star-typical CN3 branches toward extraocular muscles. **D:** Barplots showing the percentages of zebrafish exhibiting wild-type-like (“WT-like”) or defasciculated CN3 nerve(s), as scored under the dissecting stereomicroscope in G0 zebrafish at 72 hpf. Total numbers (n’s) of fish in each group are given above the corresponding bar. Data are shown from three experimental replicates. Pearson’s 2x3 chi-squared test with 2 degrees of freedom; for each of three experimental replicates, Χ^2^=129.19, p<2.2e^-16^; Χ^2^=74.61, p<2.2e^-16^; Χ^2^=52.5, p=4.0e^-12^. **E:** Kaplan-Meier survival curves demonstrating the relative survival probabilities of Scramble-injected (blue line) and Guide-injected (pink line) zebrafish over the first 72 hours of life. “Number at risk” below the plot provides the counts of surviving embryos in each group taken every 24 hpf over a 72 hpf period. Relative survival probabilities of *sema3fa*-targeting and scrambled gRNA-injected embryos were compared by the log-rank test. Displayed data were derived from a single experimental replicate, but measurements were taken for three experimental replicates, all of which showed the same trend. **F-G:** Representative images from wild-type or F2 germline *sema3fa^-/-^* mutants (arrows and stars as per A-C).

In G0 *sema3fa* mosaic null mutants, CN3 nuclei were present and appeared grossly intact. However, CN3 axons had increased defasciculation in 94.8% of fish and failed to project to their target extraocular muscles (Fig. 4A-D). There was a statistically significant association between treatment group and the proportions of fish with CN3 defasciculation, but no statistically significant difference in survival (Fig. 4D-E). Under the stereomicroscope, axonal defasciculation was visible but poorly resolved and required manual focus adjustment through z-planes to visualize by eye. This was poorly captured by stereomicroscope imaging of a single z-plane (Supplementary Figure 3A). F2 germline *sema3fa^-/-^* mutants also had increased CN3 defasciculation and failed extension toward CN3 target extraocular muscles relative to *sema3fa^+/+^* fish (Fig. 4F-G).

G0 *olig2* mutants had grossly absent (71.0%) or severely malformed (22.4%) CN6 nuclei with few residual motor neurons and absence or thinning of the CN6 nerves, reminiscent of *mafba*-null mutants (Fig. 5A-I). There was a statistically significant association between treatment group and the proportions of fish with each CN6 phenotype, but no statistically significant difference in survival (Fig. 5I-J). Like G0 mutants, F2 germline *olig2^-/-^* mutants had grossly absent or malformed CN6 motor neuron nuclei with absent or thin CN6 nerves (Fig. 5K-L).

**Figure 5.**
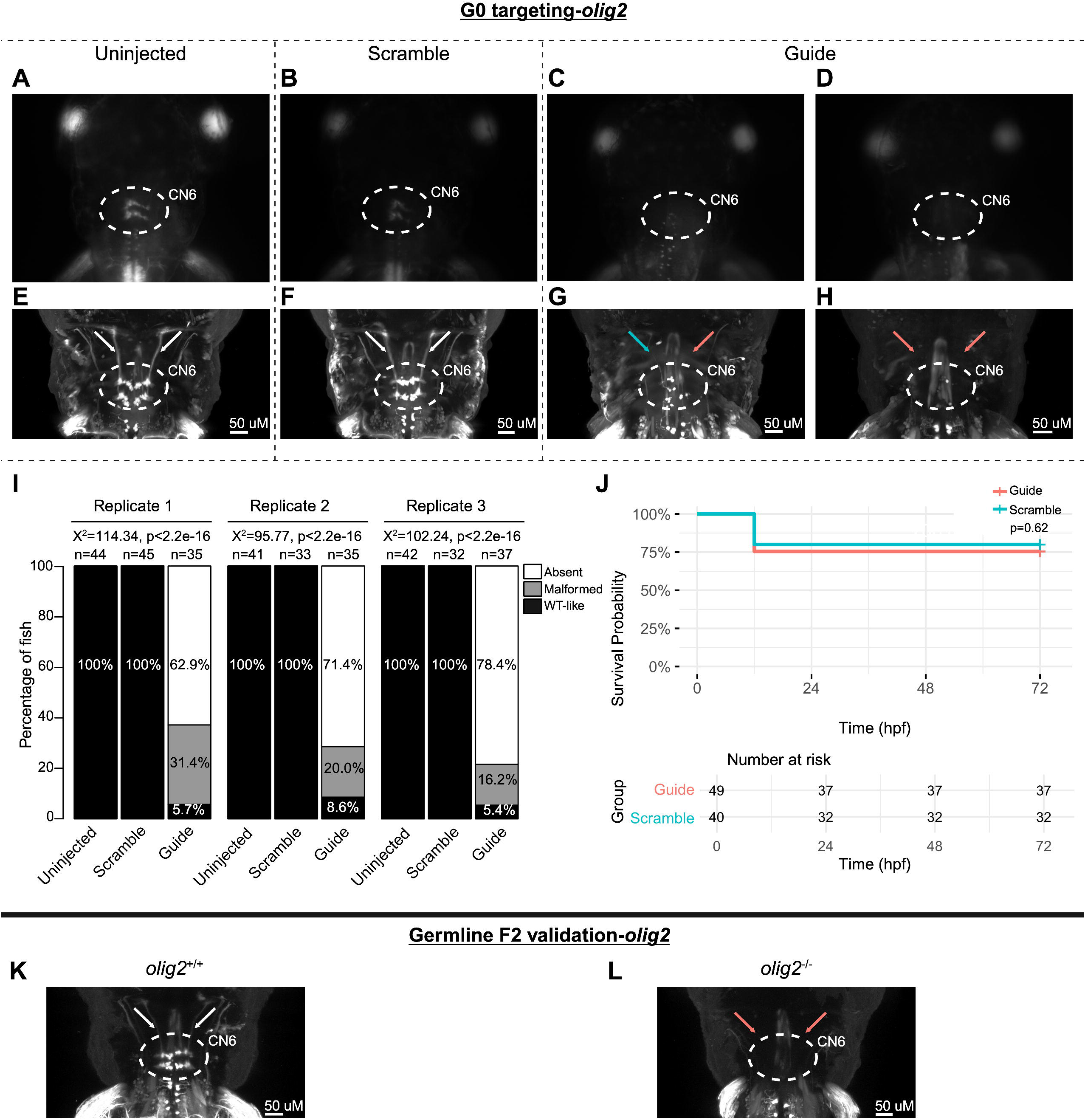
G0 mosaic and F2 germline zebrafish targeting of *olig2.* **A-H:** Initial *olig2* targeting experiments were performed in G0 embryos. Representative images obtained by stereomicroscope **(A-D)** and confocal microscopy **(E-H)** for Uninjected (A, E), Scrambled guide injected (B, F), or *olig2*-targeting Guide-injected (C-D, G-H) G0 HGj4A zebrafish at 72 hpf. Two images are provided for Guide-injected fish, to demonstrate variability in G0 targeting outcomes. Stereomicroscope images for each treatment group correspond to the confocal images obtained from the same fish in the panel below. Motor neurons in the abducens nucleus (dashed white circled region) appear absent by stereomicroscopy (C, D) or variably reduced (G, H) by confocal microscopy, and abducens nerves appear thin (G, blue arrow) or absent (G, H, red arrows) compared to normal (E, F; white arrows) by confocal microscopy. **I:** Barplots showing the percentages of zebrafish exhibiting wild-type-like (“WT-like”), malformed, or absent CN6 motor neuron nuclei, as scored under the dissecting stereomicroscope in G0 zebrafish at 72 hpf. Total numbers (n’s) of fish in each group are given above the corresponding bar. Data are shown from three experimental replicates. Pearson’s 3x3 chi-squared test with 4 degrees of freedom; Χ^2^=114.34, 95.77, 102.24 for each of three experimental replicates; p<2.2e^-16^. **J:** Kaplan-Meier survival curves demonstrating the relative survival probabilities of Scramble-injected (blue line) and Guide-injected (pink line) zebrafish over the first 72 hours of life. “Number at risk” below the plot provides counts of surviving embryos in each group taken every 24 hpf over a 72 hpf period. Relative survival probabilities of *olig2*-targeting and scrambled gRNA-injected embryos were compared by the log-rank test. Displayed data were derived from a single experimental replicate, but measurements were taken for three experimental replicates, all of which showed the same trend. **K-L:** Representative images from wild-type or F2 germline *olig2^-/-^* mutants (key as noted for A-H).

Finally, CN6 nuclei were universally present but malformed in 60.0% of *frmd4bb*-targeted G0 mutants, with reduced motor neuron dispersion through the CN6 motor nuclei. Additionally, CN6 nerves failed to reach their target lateral rectus muscles (Fig. 6A-D). There was a statistically significant association between treatment group and the proportions of fish with CN6 phenotypes, but no statistically significant difference in survival (Fig. 6D-E). F2 germline *frmd4bb^-/-^* mutants had phenotypes consistent with their G0 counterparts, including malformation of CN6 motor neuron nuclei and failed CN6 nerve extension toward target extraocular muscles relative to *frmd4bb^+/+^* fish (Fig. 6F-G). These phenotypes were also visible but poorly resolved by stereomicroscope and required manual adjustment of the focus through z-planes; thus, phenotypes were poorly captured by stereomicroscope imaging of a single z-plane (Supplementary Figure 3B).

**Figure 6.**
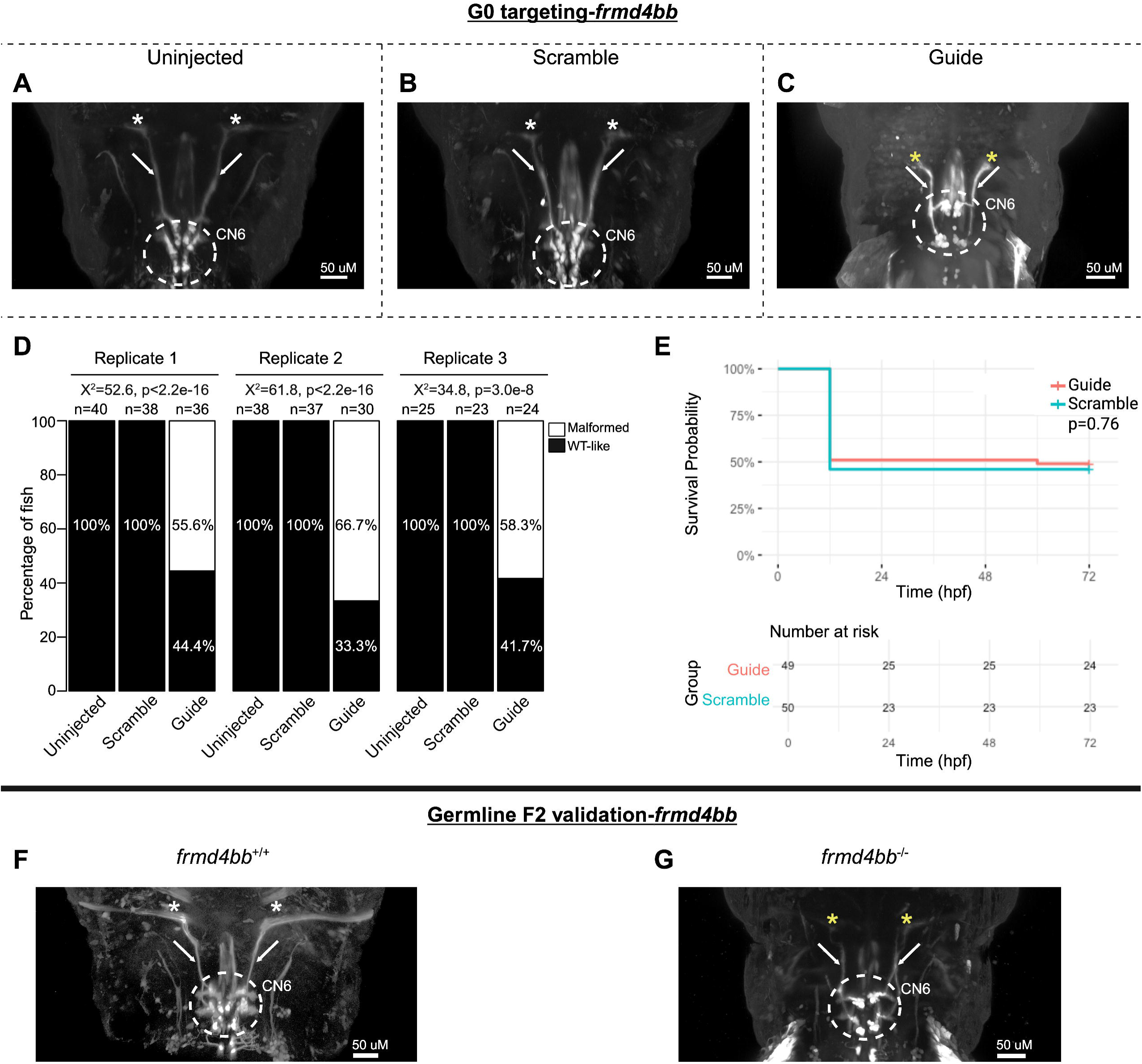
G0 mosaic and F2 germline zebrafish targeting of *frmd4bb.* **A-E:** Initial *frmd4bb* targeting experiments were performed in G0 embryos. Representative images obtained by confocal microscopy **(A-C)** of Uninjected (A), Scrambled guide injected (B), or *frmd4bb*-targeting Guide-injected (C) G0 HGj4A zebrafish at 72 hpf. Motor neurons in the abducens nucleus (dashed white circled region) have reduced dispersion through the columns of the abducens nuclei (C), and abducens nerves appear stalled and fail to target the lateral rectus muscles (C, white arrows and yellow stars) compared to Uninjected and Scrambled (A, B, white arrows and white stars). **D:** Barplots showing the percentages of zebrafish exhibiting wild-type-like (“WT-like”) or malformed CN6 motor neuron nuclei, as scored under the dissecting stereomicroscope in G0 zebrafish at 72 hpf. Total numbers (n’s) of fish in each group are given above the corresponding bar. Data are shown from three experimental replicates. Pearson’s 2x3 chi-squared test with 2 degrees of freedom; for each of three experimental replicates, Χ^2^=52.6, p<2.2e^-16^; Χ^2^=61.8, p<2.2e^-16^; Χ^2^=34.8, p=3.0e^-8^. **E:** Kaplan-Meier survival curves demonstrating the relative survival probabilities of Scramble-injected (blue line) and Guide-injected (pink line) zebrafish over the first 72 hours of life. “Number at risk” below the plot provides counts of surviving embryos in each group taken every 24 hpf over a 72 hpf period. Relative survival probabilities of *frmd4bb*-targeting and scrambled gRNA-injected embryos were compared by the log-rank test. Displayed data were derived from a single experimental replicate, but measurements were taken for three experimental replicates, all of which showed the same trend. **F-G:** Representative images from wild-type or F2 germline *frmd4bb^-/-^* mutants.

### Functional testing of transcription factor candidate variants by protein binding microarray

Our sequenced human oCCDD cohort included three pedigrees, each of which harbored a variant of uncertain significance in the DNA binding domain of one of three transcription factors. Pedigree 160 harbored c.411G>C (p.(Trp137Cys), ENST00000298231.5) in *PHOX2A*, a known CFEOM gene; Pedigree 232 harbored c.667G>A (p.(Glu223Lys), ENST00000373313.3) in *MAFB*, a known DRS gene; and Pedigree ENG_ET harbored c.467G>T (p.(Arg156Leu), ENST00000333337.3) in *OLIG2,* a novel DRS candidate gene. Interestingly, each of these genes yielded the correct cranial motor nucleus/nerve phenotype following LOF CRISPR targeting in zebrafish. To couple the gene-level testing of our zebrafish screen with functional testing of these variants of uncertain significance, we mapped the variants relative to other reported variants in these proteins (Fig. 7A-C) and assessed transcription factor-DNA interactions by protein binding microarray (Fig. 7D-L).

**Figure 7.**
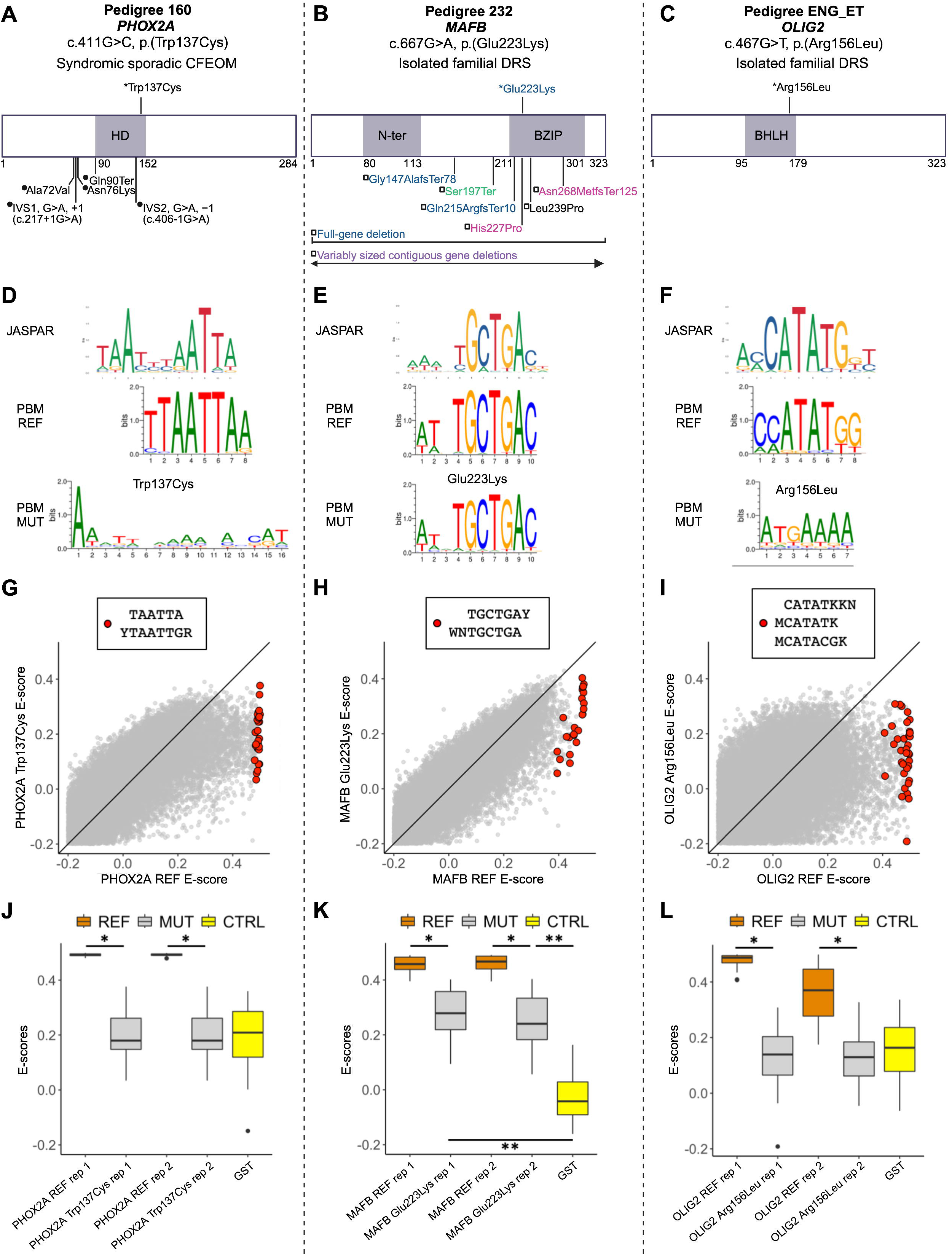
oCCDD proband-derived candidate variants in transcription factors disrupt DNA binding. **A-C:** 2D structural mapping of human variants. A: Variants in PHOX2A associated with CFEOM. B: Variants in MAFB associated with DRS and/or co-occurring phenotypes. C: Variant in OLIG2 associated with DRS. References for Figure 7 are provided in Supplementary Methods and Results. *Variants above schematics were identified and reported in our sequenced human oCCDD cohort^1^ and are functionally tested for the first time in this work: PHOX2A p.(Trp137Cys), MAFB p.(Glu223Lys), and OLIG2 p.(Arg156Leu). Variants below schematics are previously reported as follows: (A) ●-Previously reported PHOX2A variants associated with CFEOM: IVS1, G>A, +1 (c.217+1G>A);^2^ IVS2, G>A, −1(c.406-1G>A);^2,3^ p.(Ala72Val); ^2,3^ p.(Asn76Lys); ^4^ p.(Gln90Ter).^5^ (B) L-Previously reported MAFB variants associated with DRS and colored as follows: blue denotes isolated DRS;^6^ magenta denotes DRS +/-hearing impairment +/- intellectual disability;^6,7^ black denotes DRS + focal segmental glomerulosclerosis +/- hearing impairment;^8^ green denotes focal segmental glomerulosclerosis +/- DRS;^9^ purple denotes contiguous gene deletions with variable neurodevelopmental anomalies +/- DRS.^10^ Variants are mapped using the following transcripts: ENST00000298231.5 (PHOX2A), ENST00000373313.3 (MAFB), and ENST00000382357.4 (OLIG2). **D-F:** Transcription factor binding site motif logos for PHOX2A (D), MAFB (E), and OLIG2 (F). Top: logos from the JASPAR database of transcription factor binding profiles derived from high-throughput sequencing SELEX (HT-SELEX) experiments for either the wild-type human protein (PHOX2A and OLIG2) or orthologous mouse protein (Mafb). Middle and bottom: Protein binding microarray (PBM) experiment motifs derived from universal protein binding microarrays for the reference DNA binding domain (PBM REF, middle) and for the mutant DNA binding domain (PBM MUT, bottom). **G-I:** For 8-mers resembling the wild-type motif, E-score comparison between reference and mutant DNA binding domain for PHOX2A (G), MAFB (H), and OLIG2 (I). Red dots correspond to 8-mer sequences that contain the labeled International Union of Pure and Applied Chemistry (IUPAC) code-based *k*-mers, which are selected to closely resemble each motif logo. Key: Y: C or T nucleotides, R: A or G nucleotides, W: A or T nucleotides, M: A or C nucleotides, K: G or T nucleotides, N: any base. **J-L:** 8-mer E-score comparison for the *k*-mers labeled in G-I across the replicate protein binding microarrays. J: Reference versus p.(Trp137Cys) PHOX2A variant. PHOX2A p.(Trp137Cys) led to a significant drop in E-scores for 8-mers resembling the wild-type motif (*p* < 10^-9^; one-sided Mann-Whitney U test) to a level indistinguishable from E-scores of the GST negative control (*p* > 0.9; two-sided Mann-Whitney U test). K: Reference versus p.(Glu223Lys) MAFB variant. The E-scores for the 8-mers recognized by the wild-type MAFB showed significantly lower values for the mutant DNA binding domain (*p* < 10^-7^; one-sided Mann-Whitney U test), but the E-score distribution for the mutant was still higher than that of the GST negative control (*p* < 10^-6^; two-sided Mann-Whitney U test), suggesting partial loss of binding. L: Reference versus p.(Arg156Leu) OLIG2 variant. OLIG2 p.(Arg156Leu) led to a significant reduction in mutant E-scores for 8-mers recognized by wild-type OLIG2 (*p* < 10^-10^; one-sided Mann-Whitney U test) to a level similar to that of the GST-tagged negative control (*p* > 0.1; two-sided Mann-Whitney U test). Orange-reference protein, gray-mutant/ non-reference protein, yellow-GST tagged negative control. *: p<1x10^-7^; one-sided Mann-Whitney U test. **: p<1x10^-6^; two-sided Mann-Whitney U test. Abbreviations: BHLH-basic helix-loop-helix domain, BZIP-basic leucine zipper domain, CFEOM-congenital fibrosis of the extraocular muscles, CTRL-GST negative control protein, DRS-Duane retraction syndrome, GS -glutathione S-transferase tagged negative control protein, HD-homeodomain, MUT-mutant/non-reference, N-ter-N-terminal region, PBM-protein binding microarray, REF-reference (non-mutant) sequence, rep-replicate.

For each variant, we performed universal protein binding microarray experiments^4,23^ for a pair of reference (REF) and mutant (MUT) DNA binding domains on the same array and generated transcription factor binding site motif logos (Fig. 7D-F). For a quantitative view of each experiment, we analyzed the distribution of E-scores (rank-based enrichment score per DNA 8-mer ranging from -0.5 to 0.5, in which 0.5 indicates highly specific binding to that 8-mer, Fig. 7G-L).

PHOX2A p.(Trp137Cys) led to complete loss of binding to the wild-type PHOX2A motif, indicated by the degenerate motif logo for the mutant (Fig. 7D), and to a significant drop in E-scores for 8-mers resembling the wild-type motif to a level indistinguishable from that of the negative control (Fig. 7G,J).

Interestingly, MAFB p.(Glu223Lys) did not completely prevent transcription factor recognition of the wild-type motif (Fig. 7E). Nevertheless, the E-scores for the 8-mers recognized by the wild-type MAFB showed significantly lower values for the mutant DNA binding domain (Fig. 7H,K). The E-score distribution for the mutant was still higher than that of the negative control (Fig. 7K), which is indicative of partial loss of binding. This is consistent with the observation that the variant still permits the DNA binding domain to recognize the wild-type motif (Fig. 7E).

OLIG2 p.(Arg156Leu) led to a complete loss of binding to the wild-type OLIG2 motif, characterized by the degenerate mutant motif (Fig. 7F), and significant reduction in mutant E-scores for 8-mers recognized by wild-type OLIG2 to a level similar to that of the negative control (Fig. 7I,L). These data support that the tested missense variants may have a deleterious effect on protein function, but more data will be needed over time, including testing additional pathogenic and benign variants once their clinical impact has been established.

### Additional human phenotyping of pedigrees with variants in PHOX2A, MAFB, SEMA3F, OLIG2, or FRMD4B

Additional cosegregation analysis and phenotyping are shown for the human pedigrees who harbored variants in *PHOX2A, MAFB, SEMA3F, OLIG2,* and *FRMD4B* (Table 1, Fig. 8, Supplementary Table 2). The variants in these genes were absent from the population and predicted to be damaging.

**Figure 8.**
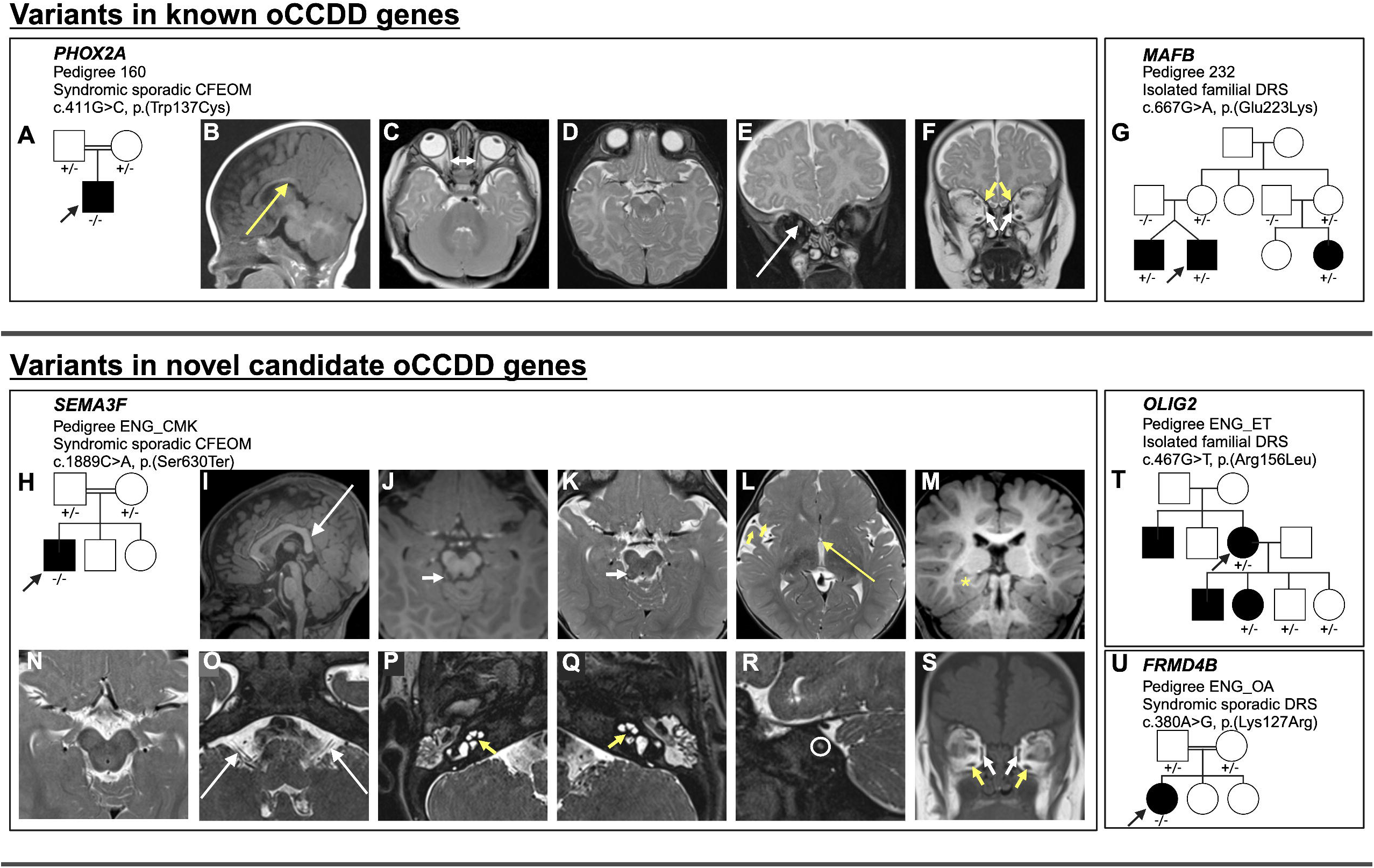
oCCDD pedigrees with functionally validated candidate genes/variants and brain MR images of probands with homozygous *PHOX2A* and *SEMA3F* variants. **A, G-H, T-U:** Schematics of pedigrees segregating variants of uncertain significance in known or novel candidate oCCDD genes. **B-F:** Images from a 1.5T Siemens brain MRI of the pedigree 160 proband at 4 months of age who has CFEOM and harbors a homozygous *PHOX2A* variant. B: Sagittal T1 fluid-attenuated inversion recovery (FLAIR) 4 mm thick image reveals abnormal anatomy of the corpus callosum with a somewhat down-slanted posterior body and splenium (long yellow arrow). C: Axial turbo spin echo (TSE) T2 weighted 4 mm thick image shows diminutive medial rectus muscles (short double-headed white arrow). D: Axial TSE T2 weighted 4 mm thick image at the level of the midbrain and interpeduncular cistern does not show oculomotor nerves. E-F: Coronal TSE T2 weighted 4 mm thick images with and without fat-suppression show asymmetric positioning of the optic nerves, higher on the right (long white arrow), diminutive medial rectus muscles (short white arrows), and small superior oblique muscles (short yellow arrows). Due to slice thickness and slice angle the superior rectus muscles were not as readily assessed but also appeared slightly small. **I-S:** The ENG_CMK proband who has CFEOM and harbors a homozygous *SEMA3F* had MR imaging at 12 months of age obtained on a 3T Siemens Skyra (I-M, O-S) and 3 months of age obtained on a 1.5T Siemens unit (N) variant. I: Midline sagittal T1 Magnetization Prepared Rapid Gradient Echo (MPRAGE) 1 mm thick image demonstrates abnormal anatomy of the corpus callosum with a somewhat down-slanted posterior body and splenium (long white arrow). J: Axial 1 mm reformatted MPRAGE and K: 2.5 mm thick TSE T2 weighted images show a small protuberance off or along the right side of the tectum (short white arrows) that appears hypointense on T1 and heterogeneous on T2 with central high signal and peripheral low signal intensity. This lesion is of uncertain etiology but was present in retrospect on an examination the previous year. L: Axial 2.5 mm thick TSE T2 weighted image demonstrating a diminutive anterior commissure (long yellow arrow) and slight underdevelopment of the right frontal and temporal opercula (short yellow arrow). M: Reformatted 1 mm coronal MPRAGE image shows asymmetry of the hippocampal formations and medial temporal lobes with the right side appearing mildly misshapen (yellow asterisk). N: Axial 2.5 mm thick T2 weighted image at the level of the midbrain does not show oculomotor nerves at the level of the interpeduncular cistern. O: Axial 0.44 mm thick T2 Sampling perfection with application optimized contrasts using different flip angle evolution (SPACE) image shows that the cisternal segments of the vestibulocochlear nerves are present but small (long white arrows) as they course posterior and parallel to the facial nerves. P-Q: T2 SPACE images through the inner ears show dysmorphic, thickened cochlear modioli with stenotic cochlear apertures, more pronounced on the left (short yellow arrows). The apices of the cochleae appear mildly flattened. R: Sagittal oblique T2 space MRI shows marked stenosis of the left internal auditory meatus (white circle) with only one cranial nerve visible instead of the expected four (facial, cochlear and superior and inferior vestibular nerves). S: Coronal 5 mm thick T1 weighted image of the brain at the level of the orbits shows small medial rectus (short white arrows) and inferior rectus muscles (short yellow arrows). The superior rectus muscles are likely also small but are suboptimally assessed due to slice thickness.

**Table 1.** Human probands’ clinical phenotypes and variants in known and novel oCCDD candidate genes derived from the zebrafish screen.

| Pedigree ID | 160 | 232 | ENG_CMK | ENG_ET | ENG_OA |
| --- | --- | --- | --- | --- | --- |
| Diagnosis | Syndromic sporadic CFEOM | Isolated familial DRS | Syndromic sporadic CFEOM | Isolated familial DRS | Syndromic sporadic DRS |
| Sex assigned at birth | Male | Male | Male | Female | Female |
| Reported previously? | PMID: 39033378 | PMID: 39033378 | PMID: 39033378 | PMID: 39033378 | PMID: 39033378 |
| **Variants** |  |  |  |  |  |
| Variant (GRCh38/hg38) | chr11:72240193C>G | chr20:40688184C>T | chr3:50186688 C>A | chr21:33027329G>T | chr3:69302379 T>C |
| Gene | *PHOX2A* | *MAFB* | *SEMA3F* | *OLIG2* | *FRMD4B* |
| Variant (cDNA, protein) | c.411G>C, p.(Trp137Cys) | c.667G>A, p.(Glu223Lys) | c.1889C>A, p.(Ser630Ter) | c.467G>T, p.(Arg156Leu) | c.380A>G, p.(Lys127Arg) |
| Transcript | ENST00000298231.5 | ENST00000373313.3 | ENST00000002829.8 | ENST00000333337.3 | ENST00000398540.8 |
| Inheritance | AR-hom | AD-inc | AR-hom | AD-inc | AR-hom |
| **Ophthalmological** |  |  |  |  |  |
| CFEOM (HP:0001491) | Y, BL | N | Y, R side | N | N |
| Congenital ptosis (HP:0007970) | Y, severe BL, L>R side | N | Y, R side | N | N |
| Impaired ocular abduction (HP:0000634) | Y, abs BL | Y, abs BL | N | Y, BL | Y, L side |
| Impaired ocular adduction (HP:0000542) | Y, abs BL | Y, abs BL | N | Y, mild BL | Y, L side |
| Limited vertical eye movements | Y, severe BL | N | Y, unable to elevate or fully depress R eye | N | N |
| Duane anomaly (HP:0009921) | N | Y, type III BL | N | Y, type I BL, upshoot L side | Y, type III with upshoot L side |
| Hypotropia (HP:0025584) | Y, R side | N | N | Y | N |
| Exotropia (HP:0000577) | Y, L side | Y | Y, intermittent R side | N | Y, in R gaze |
| Esotropia (HP:0000565) | N | N | N | Y, alternating | N |
| Ophthalmologic surgeries | N | Y, LR muscle recessions of 8 mm R side and 6 mm L side | N | Y, initial 5.5 mm MR muscle recessions BL, secondary 1.0 MR muscle recessions BL | N |
| **Other CN findings** |  |  |  |  |  |
| Absent Bell's phenomenon | Y, BL | N | N | N | N |
| CN8: Congenital sensorineural hearing impairment (HP:0008527) | N | N | Y, BL | N | ? sensorineural vs conductive R side |
| **Neurological** |  |  |  |  |  |
| Global developmental delay (HP:0001263) | N | N | Y | N | N |
| Delayed speech and language development (HP:0000750) | N | N | Y | N | Y |
| Delayed gross motor development (HP:0002194) | N | N | Y | N | Y |
| Cognitive impairment (HP:0100543) | N | N | NA | N | N |
| Attention deficit hyperactivity disorder (HP:0007018) | N | N | N | N | Y |
| Panic attacks (HP:0025269) | N | N | N | Y | N |
| Infantile muscular hypotonia (HP:0008947) | N | N | Y | N | N |
| Unsteady gait (HP:0002317) | N | N | Y | N | N |
| **Craniofacial** |  |  |  |  |  |
| Low-set, posteriorly rotated ears (HP:0000368) | N | N | Y | N | N |
| Protruding ears (HP:0000411) | N | N | Y | N | N |
| Upslanted palpebral fissures (HP:0000582) | N | N | Y | N | N |
| Wide nasal bridge (HP:0000431) | Y | N | N | N | N |
| Underdeveloped supraorbital ridge (HP:0009891) | Y | N | N | N | N |
| **Skeletal** |  |  |  |  |  |
| Overlapping toes (HP:0001845) | N | N | Y, 1st and 2nd toes BL | N | N |
| Short 2nd toe (HP:0001885) | N | N | Y, BL | N | N |
| Joint contracture of the 4th finger (HP:0009274) | N | N | Y, BL | N | N |
| Hypoplastic 5th fingernail (HP:0008398) | N | N | Y, BL | N | N |
| Short 5th finger (HP:0009237) | N | N | Y, BL | N | N |
| Volar fingernails (HP:0033976) | N | N | Y, 5th fingers and 2nd toes BL | N | N |
| Skeletal surgeries | N | N | Y, finger surgery | N | N |
| **Cardiac** |  |  |  |  |  |
| Heart palpitations (HP:0001962) | NA | NA | N | NA | Y |
| Atrial septal defect (HP:0001631) | N | NA | N | NA | Y |
| **Gastrointestinal** |  |  |  |  |  |
| Gastritis (HP:0005263) | N | N | N | N | Y |
| Gastroesophageal reflux (HP:0002020) | N | N | N | N | Y |
| Anorexia (HP:0002039) | N | N | N | N | Y |
| Decreased body weight (HP:0004325) | N | N | Y | N | N |
| **Other findings** |  |  |  |  |  |
| Cafe-au-lait spots (HP:0000957) | N | N | Y | N | N |
| Snoring (HP:0025267) | N | N | Y | N | N |
| Bony protuberance in the ear | N | N | N | N | Y, R side |

| Brain MRI |  |  |  |  |  |
| --- | --- | --- | --- | --- | --- |
| Abnl CN2<br>(HP:0000587) | Y, asymmetric positioning | NA | NA | NA | NA |
| Abnl CN3 | Y, abs BL | NA | Y, hypo BL | NA | NA |
| Abnl CN7<br>(HP:0010827) | N | NA | N | NA | NA |
| Abnl CN8<br>(HP:0009591) | N | NA | Y, cochlear nerves abs BL, vestibular nerves hypo BL | NA | NA |
| Abnl extraocular muscles<br>(HP:0008049) | Y, small MR, SO, and SR muscles BL | NA | Y, small MR and IR muscles BL, ? thin SR BL | NA | NA |
| Abnl medial temporal lobe | N | NA | Y, BL | NA | NA |
| Dysplastic corpus callosum<br>(HP:0006989) | Y, drooping splenium | NA | Y, upward bowing | NA | NA |
| Hypo anterior commissure<br>(HP:0030303) | N | NA | Y | NA | NA |
| Cochlear malformation<br>(HP:0008554) | N | NA | Y, thickened modioli, abs cochlear apertures BL | NA | NA |
| Narrow IAC<br>(HP:0011386) | N | NA | Y, BL with inferior outpouchings at the level of the fundi | NA | NA |
| Large nasopharyngeal adenoids<br>(HP:0040261) | N | NA | Y | NA | NA |
| Abnl cerebral vascular morphology<br>(HP:0100659) | Y, large vessel lateral to basilar artery | NA | N | NA | NA |

Key: ?-possibly, abnl-abnormal, abs-absent, AD-inc-autosomal dominant with incomplete penetrance, AR-hom-autosomal recessive homozygous, BL-bilateral, CFEOM-congenital fibrosis of the extraocular muscles, CN-cranial nerve, CN2-optic nerve, CN3-oculomotor nerve, CN7-facial nerve, CN8-vestibulocochlear nerve, DRS-Duane retraction syndrome, hypo-hypoplastic, IAC-internal auditory canal, IR-inferior rectus muscles, L-left, LR-lateral rectus muscles, MR-medial rectus muscles, N-no, NA-not ascertainable/ unknown, R-right, SO-superior oblique muscles, SR-superior rectus muscles, Y=yes.

The proband of syndromic sporadic Pedigree 160 harbored the homozygous *PHOX2A* variant shown above to abrogate DNA binding (c.411G>C, p.(Trp137Cys); Table 1, Fig. 8A-F). He had bilateral CFEOM with severe, bilateral restrictions in vertical eye movements; absent horizontal eye movements; severe, bilateral congenital ptosis worse on the left side; exotropia; and right hypotropia. Brain MRI revealed characteristic *PHOX2A*-associated anomalies, including indiscernible CN3 bilaterally and hypoplasia of the extraocular muscles typically innervated by CN3/CN4. Additional subtle syndromic features included mild craniofacial dysmorphisms and brain MRI findings consisting of asymmetric CN2 positioning, corpus callosal anomalies, and a prominent anomalous vessel that was likely venous in nature and coursed lateral to the basilar artery.

Three affected members of pedigree 232 and their unaffected mothers harbored a partially penetrant heterozygous *MAFB* variant shown by protein binding microarray to reduce DNA binding (c.667G>A, p.(Glu223Lys); Table 1, Fig. 8G). The family segregated three subtypes of DRS in twins and a maternal first cousin. The proband had bilateral, exotropic DRS type III characterized by bilateral limitation of abduction and adduction and globe retraction on attempted adduction. He required two eye muscle surgeries to try to improve his eye alignment; both lateral rectus muscles were recessed in each procedure. His twin had bilateral, asymmetric esotropic DRS type I with limited abduction and globe retraction on adduction bilaterally, and their cousin had left-sided exotropic DRS type II with mildly restricted adduction and globe retraction on adduction, exotropia in upgaze, myopic astigmatism, and a right head turn.

The proband of syndromic sporadic consanguineous pedigree ENG_CMK harbored a homozygous *SEMA3F* variant (c.1889C>A, p.(Ser630Ter), ENST00000002829.8) which localizes to the Ig-like domain involved in ligand binding and is predicted to result in nonsense-mediated mRNA decay (Table 1, Fig. 8H-S, Supplementary Figure 4A, Supplementary Table 5).^24^ He had syndromic right-sided CFEOM characterized by absent upgaze, limited downgaze, congenital ptosis, and intermittent exotropia. Syndromic features included bilateral sensorineural hearing impairment, mild facial dysmorphisms, and contractures and shortening of the fingers. Brain MRI revealed indiscernible CN3 bilaterally, hypoplasia of CN3-innervated extraocular muscles, small CN8, stenotic internal auditory meati, dysmorphic cochleae, corpus callosal anomalies, small anterior commissure, and asymmetric hippocampal formations and medial temporal lobes.

Two affected members of pedigree ENG_ET harbored a partially penetrant heterozygous *OLIG2* variant shown above to abolish DNA binding (c.467G>T, p.(Arg156Leu); Table 1, Fig. 8T). The proband and her daughter had DRS, and the proband’s unenrolled brother and son had DRS by report. The proband had bilateral esotropic type I DRS requiring two bilateral medial rectus muscle recessions and characterized by impaired abduction and mildly impaired adduction bilaterally, left-sided upshoot, hypotropia. The daughter of the proband had right-sided esotropic DRS type I with restricted abduction and globe retraction on adduction. Of note, two unaffected children of the proband also harbored the variant.

Finally, syndromic sporadic DRS proband ENG_OA harbored a homozygous *FRMD4B* variant (c.380A>G, p.(Lys127Arg), ENST00000398540.8) located in the Band 4.1 domain, which is involved in cytoskeletal-membrane linkage (Fig. 8U, Supplementary Figure 4B).^24^ She had left-sided exotropic DRS type III characterized by impaired abduction and adduction, globe retraction in adduction, and upshoot. Syndromic features included hearing impairment, delayed speech and walking, atrial septal defect, and gastrointestinal abnormalities.

## Discussion

Historically, linkage analysis and DNA sequencing of pedigrees segregating oCCDDs have led to the identification of recurrently mutated genes.^25–27^ More recently, our exome/genome sequencing of 467 genetically unsolved oCCDD pedigrees yielded many candidate genes/variants of uncertain significance mutated in just one pedigree.^3^ Here, our functional evaluation of a subset of these genes/variants of uncertain significance using a G0 LOF zebrafish screen and protein binding microarray further supports the oCCDD involvement of five of these.

In zebrafish, G0 and F2 LOF of known (*phox2a, mafba*) and novel (*sema3fa, olig2, frmd4bb*) oCCDD genes led to ocular cranial motor neuron developmental phenotypes. In addition, protein binding microarray testing of candidate variants in transcription factors supported the functional effects of human sequenced-derived missense alleles in known and novel candidate oCCDD genes *PHOX2A, MAFB*, and *OLIG2*.

As in homozygous LOF germline models, G0 targeting of *phox2a* resulted in loss of CN3/CN4 motor nuclei in zebrafish. Moreover, the PHOX2A p.(Trp137Cys) substitution resulted in CFEOM in a human proband and complete loss of transcription factor-DNA binding *in vitro*. Although missense alleles that localize to the DNA binding domain and predicted LOF alleles have been reported as pathogenic for *PHOX2A*-CFEOM,^1,28^ the functional effects of these alleles were not demonstrated. Thus, this is the first reported *PHOX2A*-CFEOM candidate missense allele that localizes to the DNA binding domain and is demonstrated to abolish transcription factor-DNA binding.

G0 targeting of the known DRS gene *mafba* resulted in germline homozygous null-equivalent phenotypes. Moreover, MAFB p.(Glu223Lys) resulted in reduced transcription factor-DNA binding, and the variant cosegregated with incomplete penetrance in a DRS pedigree. While reported human alleles have demonstrated reduced penetrance for DRS and other *MAFB*-associated phenotypes,^2,29–31^ functional tests of human *MAFB*-DRS alleles have thus far demonstrated haploinsufficiency or dominant negative consequences. Two DRS alleles have been reported in the MAFB DNA binding domain,^2,29,32^ but their effects on DNA binding have not been demonstrated.

*SEMA3F* encodes a semaphorin involved in axon guidance.^46^ Similar to proband ENG_CMK who had CFEOM, developmental delay, sensorineural hearing impairment, and an MRI revealing small CN3 (and inability to resolve CN4), absent CN8, and brain malformations, *Sema3f*^-/-^ mice have CN3 defasciculation and CN4 absence consistent with CFEOM, as well as hearing impairment and brain malformations.^33,34^ Although our zebrafish LOF models also had CN3 defasciculation, CN4 was grossly intact, suggesting that this phenotype may have species-specific differences (Supplementary Table 5). Interestingly, a single ClinVar submitter reported a human proband with hearing impairment and a *SEMA3F* missense variant of uncertain significance and unspecified zygosity localizing to the same Ig-like domain as the variant harbored by proband ENG_CMK (c.1849G>A, p.(Val617Met), Variation ID: 1064910; Supplementary Figure 4A), supporting putative association between *SEMA3F* variants and hearing impairment.^35^ Notably, human heterozygous *SEMA3F* missense variants of uncertain significance localizing to diverse SEMA3F protein domains were previously described in individuals with hypogonadotropic hypogonadism with or without anosmia who were not reported to have CFEOM, hearing impairment, or other syndromic features in proband ENG_CMK (Supplementary Figure 4A).^36^ Three of these reported *SEMA3F* variants were tested functionally and led to impaired protein secretion *in vitro* (p.(Thr29Met), p.(Pro452Thr), and p.(Thr724Met)). Although one variant (p.(Ala652Ser)) localized to the SEMA3F Ig-like domain, its functionality was not tested. Moreover, the proband harboring this variant also harbored a variant in *FGFR1* (p.(Arg209Cys)) which is reported as likely pathogenic for hypogonadotropic hypogonadism with or without anosmia in ClinVar (Variation ID: 548670), suggesting that this is a possible alternative explanation for the phenotype in this proband. Neither the ENG_CMK proband nor his carrier parents were noted to have anosmia or hypogonadism, but these phenotypes may have been missed in the proband due to his developmental delays and evaluation before typical pubertal onset. Alternatively, this may be attributable to the distinct nature of the proband’s homozygous nonsense variant, which is predicted to induce nonsense-mediated mRNA decay. Additional studies are required to reconcile the differences in functional consequences of variants among probands with *SEMA3F* variants.

Our G0 and F2 LOF studies and others’ published morpholino knockdown studies^37^ support that *olig2* LOF results in loss of CN6 nuclei in zebrafish, consistent with DRS. Moreover, OLIG2 is downstream of the known DRS protein MAFB,^15^ and human GWAS has associated *OLIG2* SNPs with DRS.^38^ By protein binding microarray, OLIG2 p.(Arg156Leu) resulted in a complete loss of transcription factor-DNA binding, consistent with the LOF mechanism tested by our zebrafish assay.

*FRMD4B* encodes a poorly characterized scaffolding protein involved in processes including cytoskeletal dynamics and protection against retinal dysplasia in mice.^39^ Though LOF of zebrafish *frmd4bb* results in a CN6 phenotype, *FRMD4B* is mutated in only one DRS proband in our cohort, whose specific missense variant remains untested.

In summary, because this is the first report of a *PHOX2A*-CFEOM missense variant in the DNA binding domain, pathogenicity of the p.(Trp137Cys) PHOX2A allele remains incompletely proven but is strongly suspected. Moreover, while two *MAFB*-DRS missense variants in the DNA binding domain have been reported, reduced DNA binding has not yet been demonstrated as a pathogenic mechanism. Finally, since *SEMA3F*, *OLIG2*, and *FRMD4B* are each mutated in a single pedigree in our cohort, additional human alleles resulting in a similar phenotype are needed to prove pathogenicity. However, these are now strong oCCDD candidate genes.

This G0 screening method has limitations. First, since phenotypes were screened at 72 hpf under the stereomicroscope, later-onset or less robust phenotypes may have been missed. Since many positive hits from our screen were transcription factors with obvious LOF phenotypes, focusing future screens on conserved transcription factors with known cranial motor neuron expression may lead to improved success rates. Second, our screen may not have captured LOF of all genes tested, since phenotype induction depends on inter-species conservation, mutant viability, and guide RNA and injection efficiencies (guide RNAs were not prevalidated), and because knockout of both paralogs of duplicated genes may be needed for phenotypic manifestation. Moreover, because LOF was not confirmed at the mRNA or protein levels, some may have resulted in partial LOF, resulting in incomplete phenotype manifestation. Third, the screen only examines the outcome of biallelic LOF of the tested gene; by contrast, many of the human variants were monoallelic and/or missense, and may act through alternative non-LOF mechanisms. Finally, since zebrafish lack eyelids and the levator palpebrae superioris muscle that elevates them, we were able to assess three ptosis and four MGJWS candidate genes for gross CN3 and/or CN5 abnormalities, but not for ptosis-specific CN3 vulnerabilities. Moreover, testing of DRS candidate genes in the HGj4A line enabled assessment of primary CN6 abnormalities, but not of the additional CN3 misinnervation characteristic of human DRS. For these collective reasons, we cannot definitively exclude candidates that screened negative.

Despite these limitations, our study has identified three strong novel oCCDD candidate genes and has demonstrated functionality of candidate variants in transcription factors. Our findings support that human sequence analysis can be coupled with G0 LOF screening in zebrafish and targeted functional assays to aid in the prioritization of candidate genes.

## Supporting information

Supplementary Methods and Results

## Commercial Relationships Disclosure

**J.A. Jurgens**, None; **P.M. Matos Ruiz**, None; **J. King**, None; **Emma E. Foster**, None; **L. Berube**, None; **W-M. Chan**, None; **B.J. Barry**, None; **R. Jeong**, None; **E. Rothman**, None; **M.C. Whitman**, None; **S. MacKinnon**, None; **C. Rivera-Quiles**, None; **B.M. Pratt**, None; **T. Easterbrook**, None; **F.M. Mensching**, None; **S.A. Di Gioia**, Regeneron Pharmaceutical (E); **L. Pais**, None; **E.M. England**, None; **T. de Berardinis**, None; **A. Magli**, None; **F. Koc**, None; **K. Asakawa**, None; **K. Kawakami**, None; **A. O’Donnell-Luria**, Pacific Biosciences (F), Addition Therapeutics (C,F), Tome Biosciences (C,F), Ono Pharma USA (C,F); **D.G. Hunter**, Rebion (O,P); **C.D. Robson**, None; **M.L. Bulyk**, M.L.B. is a co-inventor on U.S. patents # 6,548,021 and #8,530,638 on PBM technology and corresponding universal sequence designs, respectively (P). Universal PBM array designs used in this study are available via a Materials Transfer Agreement with The Brigham & Women’s Hospital, Inc.; **E.C. Engle**, None

## Acknowledgements

The authors thank Drs. David Schoppik and Paige Leary for sharing their invaluable expertise in zebrafish imaging.

## Supplementary Material

**Supplementary Datafile 1.** Supplementary Methods and Results Five additional supplementary tables accompany the manuscript. These have been uploaded as a separate combined supplementary spreadsheet that contains:

**Supplementary Table 1.** Human sequence-derived candidate genes and variants and their conservation in fish.

**Supplementary Table 2.** Miscellaneous phenotypic and genetic details of human pedigrees.

**Supplementary Table 3.** Gene-targeting and scrambled guide RNAs, sequencing primers, and genomic sequences targeted with zebrafish CRISPR screen.

**Supplementary Table 4.** G0 mosaic versus F2 germline mutant phenotypes for novel oCCDD candidate genes in zebrafish at 72 hpf.

**Supplementary Table 5.** Comparison of phenotypes in human, zebrafish, and mouse models of SEMA3F missense or loss of function variants.

