## Supplementary Methods and Results for "Gene identification for ocular congenital cranial motor neuron disorders using human sequencing, zebrafish screening, and protein binding microarrays"

### Index of Supplementary Methods, Results, and Figures

**Five additional supplementary tables accompany the manuscript. These have been uploaded as a separate supplementary spreadsheet:**
Supplementary Table 1. Human sequence-derived candidate genes and variants and their conservation in fish.
Supplementary Table 2. Miscellaneous phenotypic and genetic details of human pedigrees.
Supplementary Table 3. Gene-targeting and scrambled guide RNAs, sequencing primers, and genomic sequences targeted with zebrafish CRISPR screen.
Supplementary Table 4. G0 mosaic versus F2 germline mutant phenotypes for novel oCCDD candidate genes in zebrafish at 72 hpf.
Supplementary Table 5. Comparison of phenotypes in human, zebrafish, and mouse models of SEMA3F missense or loss of function variants.

#### SUPPLEMENTARY METHODS

*Prioritization of human sequence-derived alleles for screening in zebrafish*

In our previous study, we leveraged human genetics to prioritize novel oCCDD candidate genes/ variants through phenotyping and exome/ genome sequencing of a large cohort of human pedigrees with oCCDDs.[1] Candidate genes/ variants were prioritized through multiple modalities including allele frequency filtering, predictive scores, pedigree-based analyses, identification of recurrently mutated genes and recurrent variants, annotation of animal models in the Monarch database, *de novo* variant analyses, and gene ontology analyses. In the present study, we further prioritized these candidate genes and variants based on amino acid-level conservation between human and zebrafish, recessive inheritance, and/or putative loss-of-function (LOF) consequences of the candidate variants (defined as stopgain, stoploss, frameshift, or splice site variants). Several human pedigrees had additional novel candidate genes/ variants which were not highlighted in our previous study, but which were also prioritized for the zebrafish screen based on conservation in fish, published literature, *in silico* predictors, and/or putative LOF consequences of the variant.

*Sanger validation and cosegregation analysis of human sequence-derived variants*

Sanger sequence validation was performed for candidate variants in genes that 1) yielded cranial motor phenotypes in both G0 and F2 mutants in our zebrafish screen, and/or 2) encoded transcription factors that were prioritized for protein binding microarray testing. PCR primers were designed using Primer3 v4.1.0[2] and assessed for specificity of amplification relative to other sites in the human genome (build GRCh38) using BLAT.[3] Sanger sequencing was performed for validation of variants in the probands and cosegregation analysis in additional pedigree members, when available (Supplementary Table 2).

*Guide RNA design and synthesis*

Using CHOPCHOP[4–6] (accessed November 2018 (v2) and February 2020 (v3)), the eight highest-ranked guide RNAs (gRNAs) per target gene without predicted protein-coding off-target sites were selected. gRNAs that were also represented in a published 4-guide lookup table were prioritized.[7] When possible, gRNAs targeting coding sequences at least 50 bp upstream of the penultimate exon-exon junction were selected to induce nonsense-mediated mRNA decay.[8] The top four remaining guides with the fewest predicted off-target effects and highest predicted efficiencies were selected, and PAM sequences were omitted. Each of the four guides was submitted to the Genscript scrambled sequence generator to identify non-targeting scrambled guide control sequences. The first 2 nucleotides of gene-targeting and scrambled guides were modified to start with "GG" and flanked with a 5' and 3' sequence (TAATACGACTCACTATA; GTTTTAGAGCTAGAAATAGC) to generate top-strand oligos for annealing (Supplementary Table 3). The four top-strand gene-targeting or scrambled control oligos per gene were pooled (final pooled concentration 0.2 uM) and annealed with a universal bottom-strand ultramer (Integrated DNA Technologies, final concentration 0.2 uM) as described.[9] *In vitro* transcription was performed using the MEGAscript T7 Transcription Kit (Thermo Fisher Scientific, Cat #AM1334) following the manufacturer's protocol, except for overnight incubation (≤16 hours) to increase RNA yield. RNA was purified using RNA Clean and Concentrator-5 kit (Zymo, Cat #R1016) following the manufacturer's protocol, except for use of an 8 uL elution volume to concentrate RNA.

*Guide RNA/Cas9 complex formation and microinjection*

Alt-R S.p. Cas9 Nuclease V3 (Integrated DNA Technologies, Cat #1081059) was diluted to yield a 10 uM Cas9 solution in 20 mM Tris-HCl, 600 mM KCl, and 20% glycerol and stored at -20°C. Cas9 solution and gRNAs were mixed for a final concentration of 5 uM Cas9, 1 ug/uL gRNA. When guides were pooled, four gRNAs were mixed such that total gRNA concentration remained 1 ug/uL. Cas9/gRNA mixture was incubated at 37° C for 5 minutes to generate Cas9/RNP complexes. The yolks of single-cell stage embryos were microinjected with 0.5-1.0 nL of Cas9/gRNA mixture.

*Zebrafish husbandry*

Zebrafish experiments were approved by the BCH Institutional Animal Care and Use Committee, and standard fish care was performed by the BCH Aquatic Resources Facility. Zebrafish were maintained on a standard 14 hour light/10 hour dark cycle at 28.5°C. Before being added to the system at 5 dpf, embryos and larvae were maintained in 10 cm dishes with 30 mL of sterile fish water and densities of 30-50 fish per dish. For embryos younger than 24 hpf, water was supplemented with 0.5 ppm methylene blue. To avoid long-term isolation, individually genotyped adult fish were tagged with identifiable visible implant elastomers and housed in groups (Northwest Marine Technology, 2017).[10]

*G0 screening and F2 germline validation in LOF zebrafish models of prioritized genes*

Experiments targeting CFEOM, ptosis, and MGJWS or DRS candidate genes were conducted in *Tg(isl1:GFP)*[11] or HGj4A *mnr2b/hlxb9lb*[12] reporter fish, respectively. The *Tg(isl1:GFP)* reporter line was originally generated through transgenic introduction of a linearized GFP-tagged *islet1* promoter/enhancer sequence, and the HGj4A line was made by *Tol2* transposition-mediated enhancer trapping to introduce a GFP construct upstream of the *mnr2b/hlxb9lb* gene.

Zebrafish embryos were generated using timed incrosses of adult reporter fish. G0 targeting experiments consisted of microinjecting single cell-stage embryos with four high dose (1ug/uL) guide RNAs redundantly targeting each gene.[7] Following injections, dead embryos and debris were removed twice daily. Live embryos were incubated at 28.5°C and counted every 24 hours until 72 hours post-fertilization (hpf). Sterile fish water with 0.2 mM N-

Phenylthiourea, ≥98% (Sigma-Aldrich, Cat #P7629-100G) was added at 24 hpf to inhibit pigmentation/ melanization and replaced every 24 hours.

At 72 hpf, injected G0 fish were assessed using the Nikon SMZ1500 fluorescent stereomicroscope and NIS Elements AR 5.21.03 software to assess for gross phenotypic changes in cranial motor neuron nuclei and/or nerves. To visualize abnormalities at multiple z plane levels within each fish, we manually adjusted the focus level of the stereomicroscope through multiple z planes that collectively encapsulated the anatomic regions of interest. Fish were assessed for absent or malformed motor nuclei and aberrant axonal projections of CN3, CN4, and CN5 (*Tg(isl1:GFP)* fish) or CN6 (HGj4A fish). Preliminary G0 fish phenotypes were derived from single experimental replicates without detailed phenotyping. Two additional G0 experimental replicates and F2 germline mutant validations were performed for genes whose targeting induced putative cranial motor nucleus/ nerve phenotypes in at least a subset of injected fish; these additional experiments were performed with confocal imaging.

*Zebrafish confocal image acquisition and processing*

G0 fish with putative cranial nucleus/ nerve phenotypes visualized under the stereomicroscope and F2 germline fish were additionally phenotyped using confocal imaging. G0 mutants were phenotyped blindly relative to wild-type uninjected or scrambled guide RNA-injected clutchmates. Imaging was performed on fish from at least three independent clutches for both G0 and F2 experiments. At 72 hpf, zebrafish were anesthetized and mounted dorsally in 1% low melting point agarose (ThermoFisher Scientific, Cat #16520100) in fish water in FluoroDishes (World Precision Instruments, Cat #FD3510). Fish were live imaged with a Zeiss LSM980 series upright laser scanning confocal microscope with a 20X water dipping objective (Cat #421452-9800-000). Images were acquired using Zen Software (Carl Zeiss MicroImaging GmbH, Göttingen, Germany) with 1024x1024 pixels, scan speed of 5, and 1 μM z-stacks.

Three-dimensional confocal z-stack images were processed using Arivis Vision4D software v4.0. The purpose of image processing was to improve standardization of experimentally-matched images and, when necessary, to remove non-cranial motor neuron/nerve anatomic structures that would otherwise obscure the pertinent anatomic features highlighted within each mutant. Image processing consisted of: 1) rotating images so that key anatomic structures were captured in a standard manner within a single 2-dimensional X-Y plane of control and mutant fish imaged in the same experiment; 2) transforming pixels back to their original dimensions for each image; 3) cropping and setting equivalent zoom levels for experimentally-matched images so that the same anatomic structures were encompassed within each; 4) creating a standard-sized scale bar in the 2D z-plane in which a standard anatomic landmark was present for all experimentally-matched images; 5) digitally masking extreme autofluorescence from the zebrafish eyes and skin, which would otherwise obscure the cranial motor neuron/ nerve anatomy of interest; 6) for sema3fa images, manual cropping of non-CN3 anatomic structures to show only the pertinent anatomy in processed images; 7) generating a high-resolution 3D rendering of the final image. Raw unprocessed confocal images are provided for G0 and F2 fish (Supplementary Figures 1 and 2).

*Protein structural mapping and universal protein binding microarray testing of transcription factor candidate variants*

2D protein structural maps were generated based on domain annotations in InterPro v101.0.[13] As described previously,[14] protein binding microarrays were used to assess DNA binding capabilities of variants of uncertain significance in the DNA binding domains of known (*PHOX2A, MAFB*) or novel (*OLIG2*) transcription factor-encoding candidate genes relative to their wild-type counterparts.

For protein binding microarray experiments, gBlock Gene Fragments encoding the DNA binding domains of human PHOX2A, MAFB, and OLIG2 were synthesized as double-stranded DNA fragments purchased from Integrated DNA Technologies. These fragments were cloned into a Gateway cloning-compatible entry vector, pDONR-221 (Invitrogen, Cat#12536017). Mutations PHOX2A-p.(Trp137Cys), MAFB-p.(Glu223Lys), and OLIG2-p.(Arg156Leu) identified from human oCCDD exome or genome sequences were introduced into the vectors using QuickChange site-directed mutagenesis (Agilent, Cat#200519) and mutagenic PCR primers optimized according to the QuickChange primer design manual. Sequence verification was performed using Sanger sequencing through the Harvard Medical School Biopolymers Facility to confirm correct mutagenesis. Subsequently, wild-type and mutant constructs were cloned into N-terminal GST-tagged Gateway-compatible pDEST15 vectors (Invitrogen, Cat#11802014) and confirmed again by Sanger sequencing.

Wild-type and oCCDD mutant transcription factor DNA binding domain proteins were expressed using PURExpress in vitro transcription/translation Protein Synthesizer Kits (NEB, Cat#E6800L) in the same batch. Protein expression and correct size were validated by Western blot using rabbit anti-GST primary antibody (Sigma, Cat#G7781) and goat anti-rabbit secondary antibody (Pierce, Cat#31460). Protein concentration was quantified from the Western blots using GST protein standards (Pierce, Cat#20237) and analyzed with ImageJ software.[15]

Protein binding microarrays were prepared by following a standard double-stranding primer extension reaction with ThermoSequenase DNA Polymerase (Cytiva, Cat#E790000Y), unlabeled dNTPs (NEB, Cat#N0447S), and fluorescently labeled Cy3-dUTPs (Cytiva, Cat#PA53022). Arrays were then scanned for Cy3 signal at 523 nm with a 500 lp filter and analyzed using the Double Stranding Analysis pipeline. Wild-type and oCCDD mutant DNA binding domains were assessed using a custom-designed Universal Protein Binding Microarray (8 x 60K GSE format, AMADID #030236; Agilent Technologies, Inc.) in phosphate-buffered saline (PBS) buffer.

Wild-type and mutant PHOX2A proteins were assayed at a concentration of 400nM on the PBM. Wild-type and mutant MAFB and OLIG2 proteins were assayed at a concentration of 600nM on the PBM. Each wild-type DNA binding domain and its corresponding mutant were tested in a separate chamber on the same array. Following protein binding, arrays were incubated with Alexa Fluor 488-conjugated anti-GST antibody (Invitrogen, Cat#A-11131) for fluorescent detection. Protein binding microarrays were then scanned using a GenePix 4400A microarray scanner. GPR files were generated using GenePix Pro 7.0 software at settings of 500, 400, and 300 PTM. Analysis of GPR files was performed using the Bulyk Lab PBM Universal Analysis Suite.

*Figure generation*

Figure schematics were created with BioRender.com using an academic license through Boston Children’s Hospital. Licenses for main figures: AD27AKZL6I, KC27AL00ZM, ED27AL086K, HC27AL0GI6, LU27AL0PWA, AN27AL0WI1, ZY27AL149W, AG27ANNN5T. Licenses for supplementary figures: *UJ276VPQK0, GI276VPOCW, YV27819I1A, MT277FW9XW.*

#### SUPPLEMENTARY FIGURES

*Supplementary Figure 1. Unprocessed G0 confocal images*

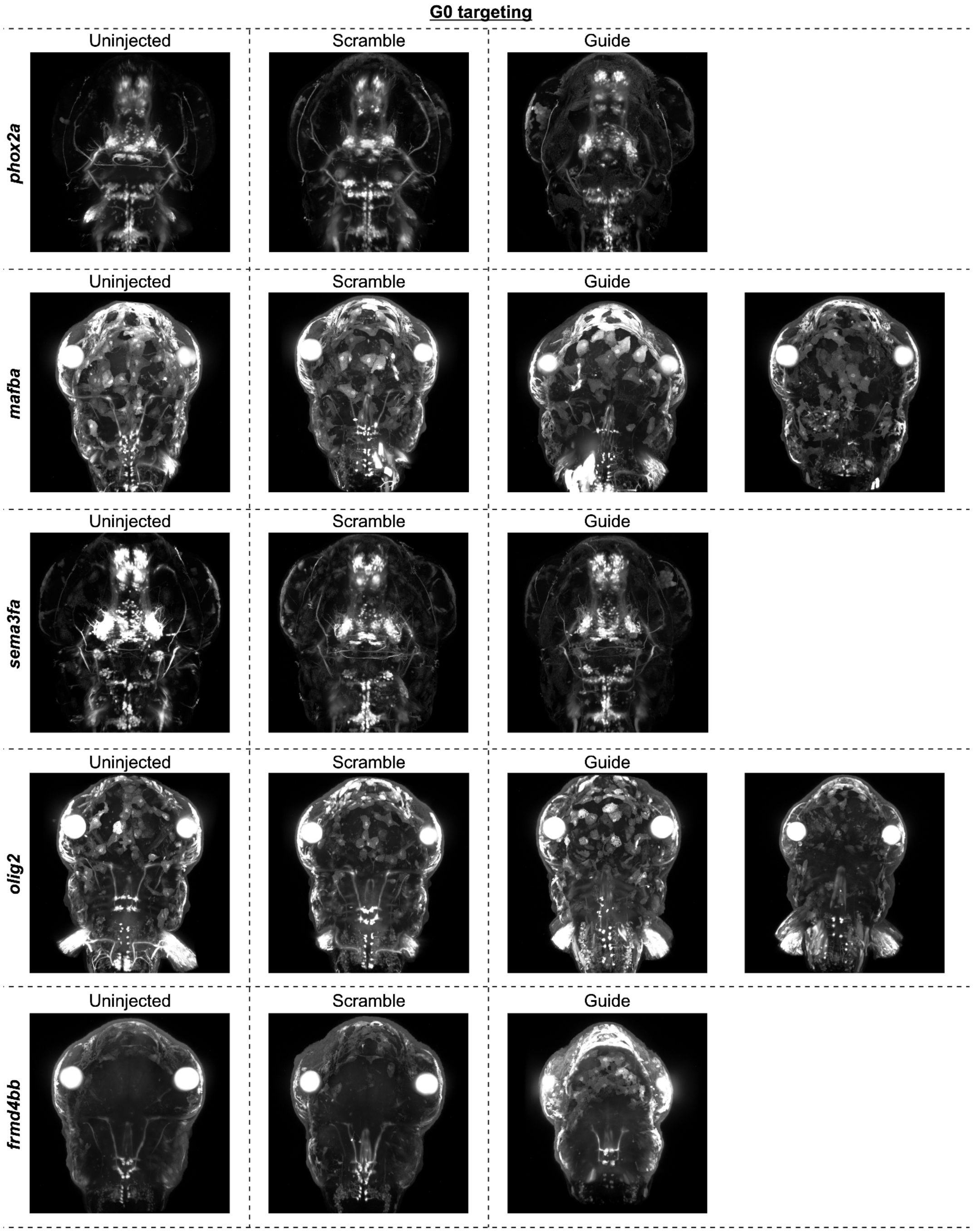

For each tested gene (named on the far left side of the figure), uninjected (left), scramble guide-injected (middle), and gene-targeting guide-injected (right) zebrafish on the appropriate reporter line backgrounds were imaged using confocal microscopy. Unprocessed confocal images are shown here and demonstrate autofluorescence of the skin and eyes, which in some cases obscure anatomical structures of interest. While non-ocular abnormalities were observed in some mutants (e.g., malformation of spinal and vagus motor neurons in *mafba* and *olig2* mutants), these were not the primary phenotypes assessed by our screen.

*Supplementary Figure 2. Unprocessed F2 confocal images*

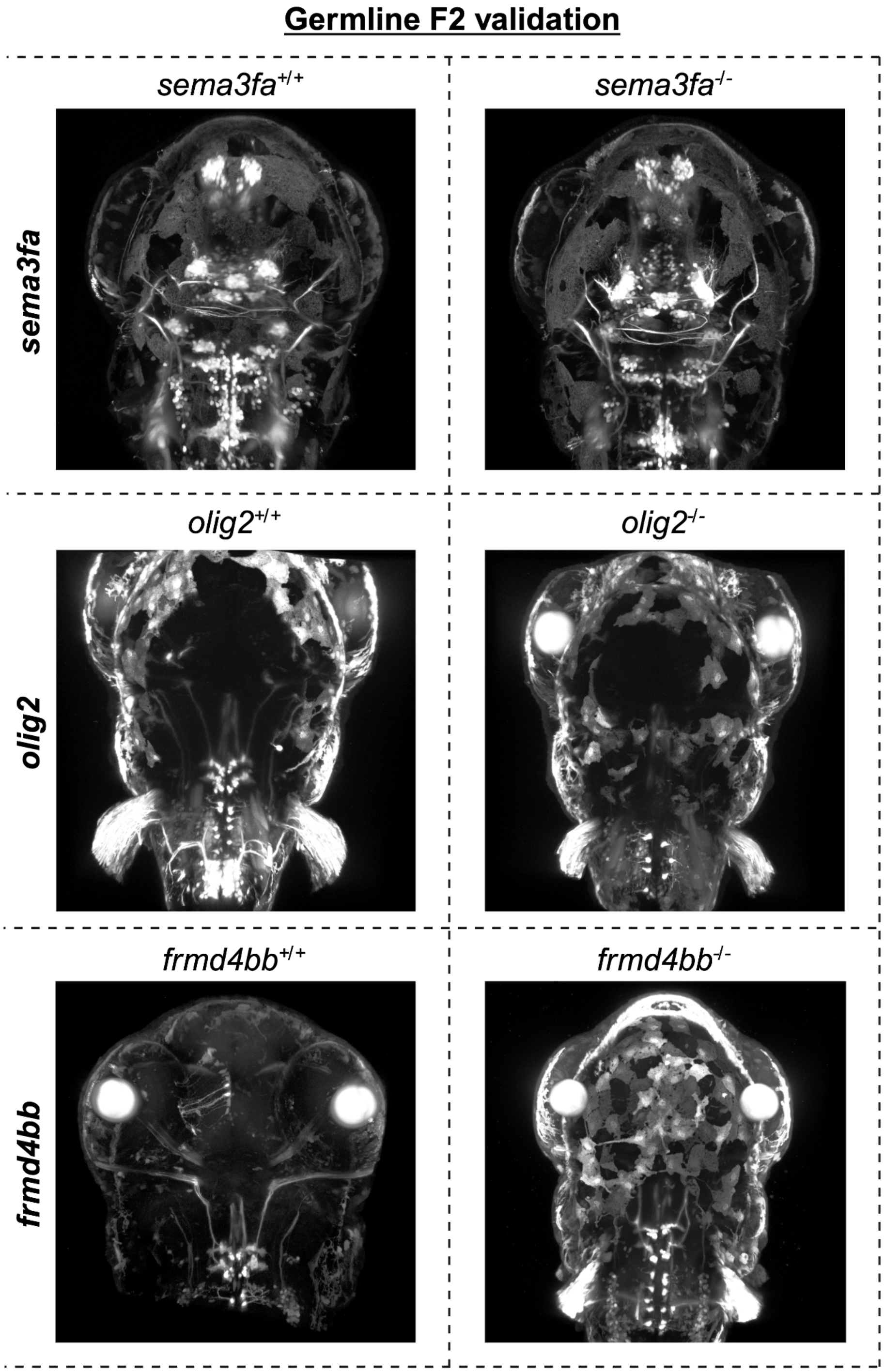

For each tested gene (named on the far left side of the figure), wild-type (left) and homozygous null (right) mutant zebrafish on the appropriate reporter line backgrounds were imaged using confocal microscopy. Unprocessed confocal images are shown here and demonstrate autofluorescence of the skin and eyes, which in some cases obscure anatomical structures of interest. While non-ocular abnormalities were observed in some mutants (e.g., malformation of spinal and vagus motor neurons in *mafba* and *olig2* mutants), these were not the primary phenotypes assessed by our screen.

*Supplementary Figure 3. Stereomicroscope images of sema3fa- and frmd4bb-targeted G0 fish*

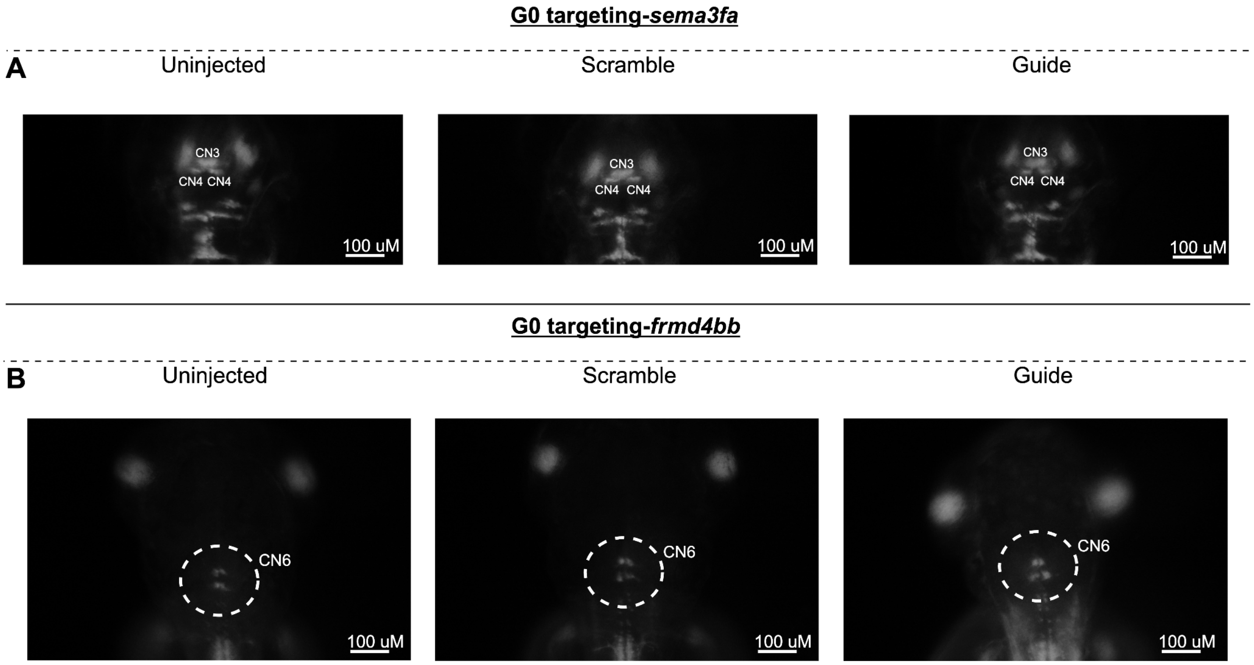

For each tested gene (named on the far left side of the figure), uninjected (left), scramble guide-injected (middle), and gene-targeting guide-injected (right) zebrafish on the appropriate reporter line backgrounds were imaged in a single z-plane on the stereomicroscope. While phenotypes including axonal defasciculation (*sema3fa*) or condensation of the columns of the motor nuclei and shortened nerves (*frmd4bb*) were visible under the stereomicroscope, these appeared more subtle than the obvious motor neuron loss phenotypes and required manual adjustment of the focus up and down in z-space to visualize. For this reason, confocal imaging was required to encapsulate the phenotypes of these mutants.

*Supplementary Figure 4. 2D protein structural mapping of SEMA3F and FRMD4B*

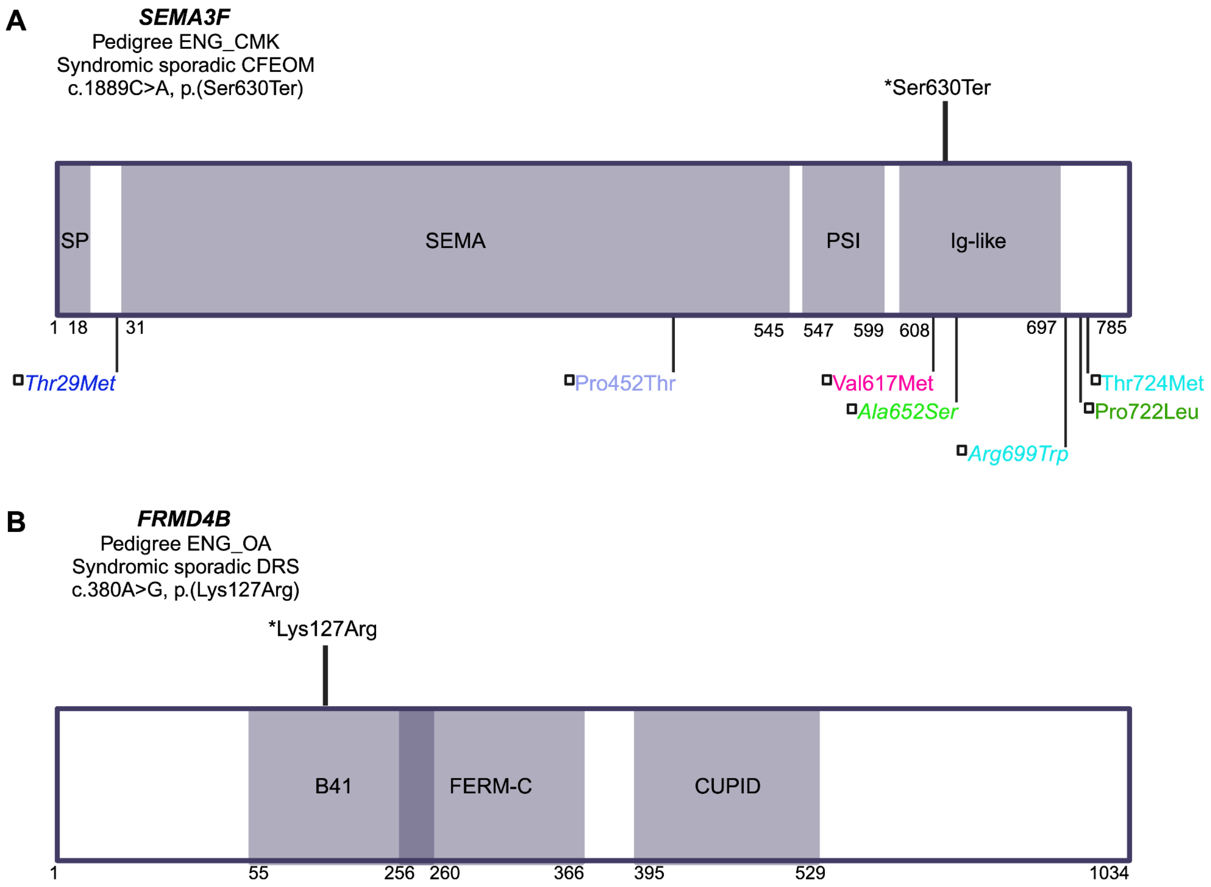

2D structural mapping of human variants. A: Variants in SEMA3F associated with CFEOM and/or other SEMA3F-associated phenotypes. B: Variants in FRMD4B associated with DRS. Key: *Variants above schematics and colored in black were initially identified and reported in our sequenced human oCCDD cohort[1] and LOF of these genes is tested for the first time in zebrafish in this work: **A)** *SEMA3F* c.1889C>A, p.(Ser630Ter) with CFEOM + sensorineural hearing loss + developmental delay + brain malformations + mild facial dysmorphisms + shortening and contractures of the fingers, and **B)** *FRMD4B* c.380A>G, p.(Lys127Arg) with DRS + hearing impairment + delayed speech and walking + atrial septal defect + gastrointestinal abnormalities. **A)** ☐-Variants below schematic are previously reported heterozygous SEMA3F missense variants, colored as follows: dark blue denotes reproductive system phenotype + anosmia/ hyposmia + obese/ overweight; light blue denotes +/- reproductive system phenotype +/- anosmia/ hyposmia +/- obese/ overweight; lime green denotes reproductive system phenotype + anosmia/ hyposmia; cyan denotes reproductive system phenotype; dark green denotes reproductive system phenotype + obese/overweight.[16] Note that three italicized variants from this reported SEMA3F case series (SEMA3F p.(Thr29Met), p.(Ala652Ser), and p.(Arg699Trp)) also carried variants of uncertain significance in additional genes known to be associated with the probands' conditions *(CHD7* and *IGSF10; FGFR1;* and *TACR3,* respectively). The probands' specific alleles in each of these three genes had each been reported in the literature in at least one additional unrelated proband with similar phenotypes. Magenta denotes hearing impairment (ClinVar Variation ID: 1064910).[17] Variants are mapped using the following transcripts: ENST00000002829.8 (SEMA3F), ENST00000398540.8 (FRMD4B). Abbreviations: B41- band 4.1 domain, CFEOM- congenital fibrosis of the extraocular muscles, CUPID- cytohesin ubiquitin protein inducing domain, DRS- Duane retraction syndrome, FERM-C- FERM C-terminal PH-like domain, Ig-like- immunoglobulin-like domain, PSI- plexin semaphorin integrin homology domain, SEMA- semaphorin domain, SP- signal peptide region.

**FIGURE 7 REFERENCES**
